# Armeniaspirols inhibit the AAA+ proteases ClpXP and ClpYQ leading to cell division arrest in Gram-positive bacteria

**DOI:** 10.1101/685669

**Authors:** Puneet Labana, Mark H. Dornan, Matthew Lafrenière, Tomasz L. Czarny, Eric D. Brown, John P. Pezacki, Christopher N. Boddy

## Abstract

Multi-drug resistant bacteria present an urgent threat to modern medicine, creating a desperate need for the discovery of antibiotics with new modes of action. Natural products whose unique highly diverse structures have been shaped by evolution to possess biologically relevant activity are an ideal discovery ground for new antibiotics with new mechanisms of action. In this study we elucidate the mechanism of action of the Gram-positive antibiotic armeniaspirol, a compound for which resistant bacteria could not be selected for. We show that armeniaspirol inhibits the ATP-dependent proteases ClpXP and ClpYQ in biochemical assays and in the Gram-positive bacteria Bacillus subtilis. We then show that this activity dysregulates key proteins involved in the divisome and elongasome including FtsZ, DivIVA, and MreB all of which are known to inhibit cell division when upregulated. Inhibition of ClpXP and ClpYQ leading to dysregulation of the divisome and elongasome represents a new mechanism of action and armeniaspirol is the first known natural product inhibitor of the coveted anti-virulence target ClpP. Thus armeniaspirol is the lead compound for a promising new class of antibiotics with a unique pharmacology and a novel mechanism for combating antimicrobial resistance, making it a highly promising candidate for further development.

## Introduction

The emergence of multi-drug resistant bacterial pathogens threatens to upend the modern era of antibiotics. The majority of clinical antibiotics inhibit a small number of cellular targets including the bacterial cell wall, ribosomes, or DNA^1^. These similar mechanistic targets enable high-level multi-drug resistance mechanisms to be readily transferred between bacteria. As a result, an increasing number of pathogens are becoming resistant to many, sometimes all, of our clinical antibiotics and too few antibiotics effective against these resistant pathogens are in development. Thus, the discovery of antibiotics that interact with new biochemical targets, and for which resistant bacterial strains do not readily emerge, are urgently needed to combat antimicrobial resistance^2,3^.

Nearly all antibiotics are derived from natural products as they permit access to a unique combination of highly diverse chemistry and bioactivity shaped through evolution that is near impossible to replicate in the laboratory. Armeniaspirols A-C represent a novel class of natural product antibiotics initially isolated from *Streptomyces armeniacus*^4^ (**Figure 1a**). These structurally unprecedented compounds were shown to be active against multi-drug resistant Gram-positive bacteria including methicillin-resistant *Staphylococcus aureus* (MRSA), vancomycin-resistant *Enterococcus* (VRE) and penicillin-resistant *Streptococcus pneumoniae* (PRSP) at low micromolar concentrations^4,5^. Furthermore, armeniaspirol A is active against MRSA in an *in vivo* murine sepsis model, and significantly, *in vitro* resistant *S. aureus* strains could not be generated even after 30 serial passages under sub-lethal doses^4^. The unique scaffold and potent antibiotic activity of armeniaspirol warranted further investigation into its unknown mechanism of action.

**Figure 1.**
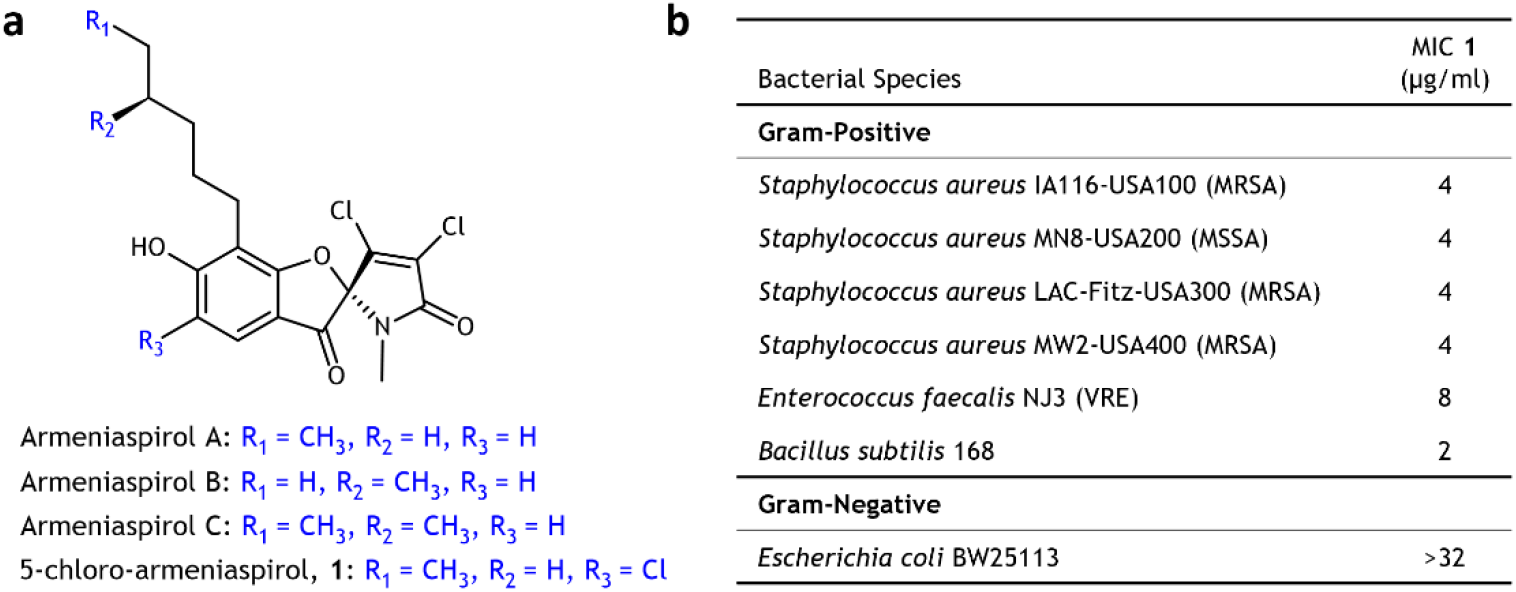
Structures of armeniaspirols. **A - C.** (a) Armeniaspirols are natural products isolated from *Streptomyces armeniacus* and possess a unique spiro-[4.4]non-8-ene scaffold. A synthetic route to armeniaspirol A generated a racemic mixture of the chlorinated derivative 5-chloro-armeniaspirol A, **1**. (b) **1** is active against Gram-positive bacteria, including pathogenic MRSA and VRE.

Using a combination of chemical synthesis, chemical and quantitative proteomics, microscopy and biochemical assays, we show that armeniaspirols directly inhibit the AAA+ (**A**TPase **A**ssociated with diverse cellular **A**ctivities) proteases ClpYQ and ClpXP. Inhibition of the ClpXP and ClpYQ proteases in *Bacillus subtilis* leads to an increase in abundance of key divisome and elongasome proteins, including FtsZ, DivIVA, and MreB, which ultimately inhibits cell division. Armeniaspirol represents the first natural product inhibitor of the promising anti-virulence target ClpP, and its dual inhibition of ClpXP and ClpYQ enables it to effect potent antibiotic activity. This novel mechanism of action and lack of detectable resistance makes armeniaspirol an exciting candidate for antibiotic development.

## Results

### Armeniaspirol does not interact with common antibiotic targets

*B. subtilis* is one of the best studied Gram-positive bacteria and serves as a model organism for Gram-positive pathogens including *S. aureus*^6^. Thus to uncover mechanism of action, we applied dimethyl isotopic labelling^7^ to quantitatively profile changes to the proteome of *B. subtilis* upon treatment of cultures with sub-lethal levels of synthetic chloro-armeniaspirol, **1** (**Figure 1a, Scheme S1**). This readily accessible synthetic derivative possesses comparable antibiotic activity to armeniaspirol A^5^, with a low minimum inhibitory concentration (MIC) against the Gram-positive pathogens MRSA and VRE (**Figure 1b**).

We profiled the impact of 1 μg/mL (2.4 μM, ½ of the MIC) of **1** on the *B. subtilis* proteome (**Figure 2a-c, Supplementary File 1**). Comparison of the proteome of the *B. subtilis* treated with **1** to *B. subtilis* treated with antibiotics that inhibit protein biosynthesis (tetracycline and chelocardin)^8^, fatty acid biosynthesis (platencin, platensimycin, cerulenin and triclosan)^9^, cell wall biosynthesis (merscadin, bacitracin, and vancomycin)^10^, and those that act as DNA damaging agents (mitomycin, daunomycin, and adriamycin)^11,12^ showed that **1** elicited a unique fingerprint. Specifically, increases in abundance of key proteins upon treatment with each of these antibiotics are not observed with **1**. Most relevant proteins showed a statistically and biologically insignificant change (**Figure 2d, Table S1**). Moreover, checkerboard synergy assays showed **1** functions independently of tetracycline and cerulenin, as well as the cell wall biosynthesis inhibitor penicillin, and the DNA replication inhibitor ciprofloxacin (all FIC indexes >0.5; **Table S2**). Together, these data indicate that armeniaspirol does not inhibit the function of common antibiotic targets and instead suggests a novel target and mechanism.

**Figure 2.**
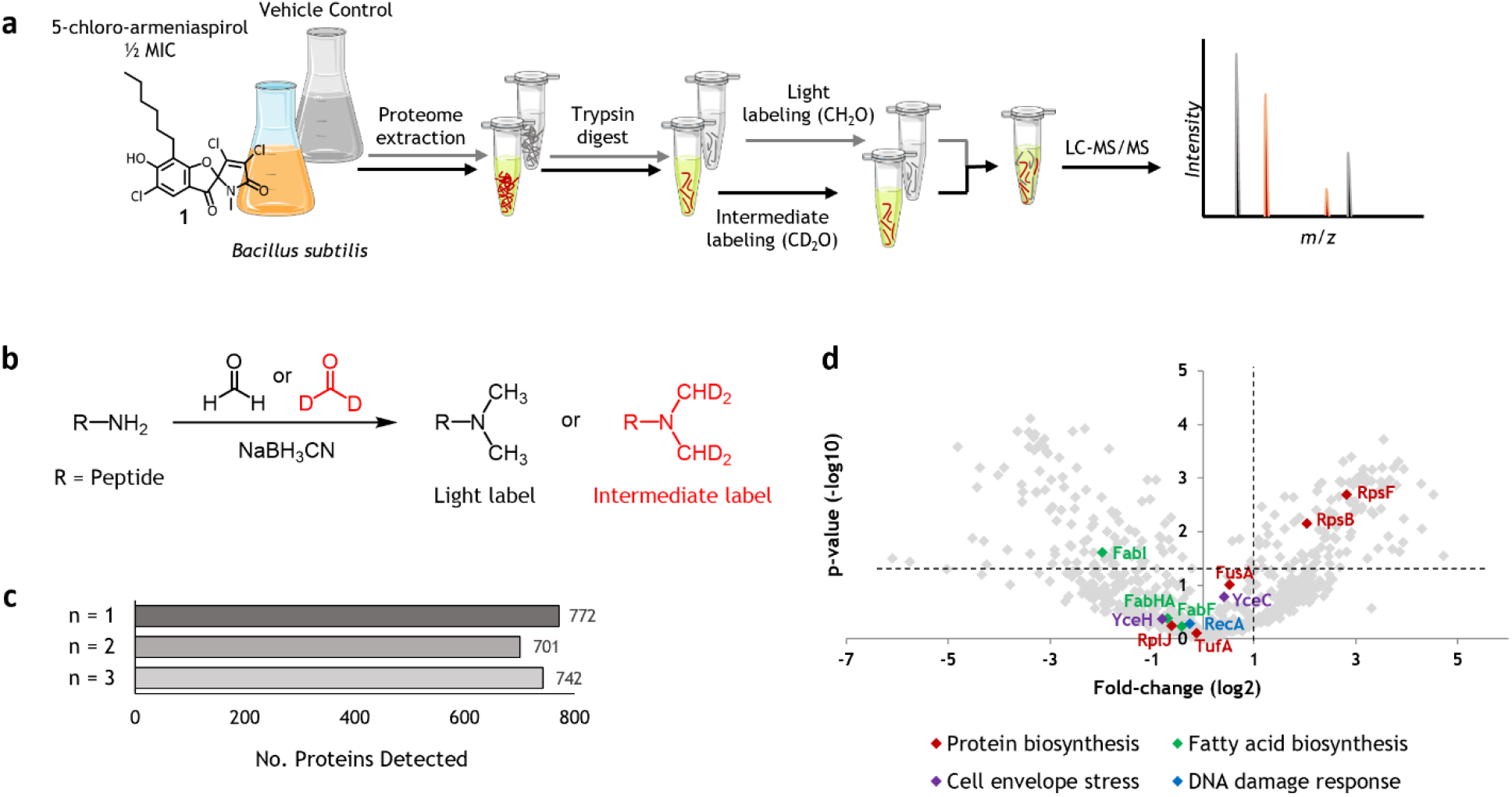
Dimethyl labeling and quantitative proteomics of *B. subtilis* proteome. (a) Proteome extract of *B. subtilis 168* from **1**- and vehicle control-treated cells are labelled with formaldehyde and deuterated formaldehyde, respectively and pooled for LC-MS/MS analysis. Experiments were carried out in biological triplicate (n=3). (b) Stable isotope dimethyl labelling chemistry. (c) Total protein recovery following dimethyl labelling for each replicate is shown. (d) Armeniaspirol-treated proteome in relation to protein biomarkers involved in various antibiotic mechanisms. Significance is defined as fold-change > 2 and p-value < 0.05. Each protein marker highlighted has been shown to have substantial enrichment with its corresponding antibiotic treatment.

### Target discovery by inhibitor capture

We thus set out to capture direct biochemical targets of armeniaspirol from the *B. subtilis* proteome. To design a capture probe, several derivatives of **1** were synthesized and their MICs determined (**Figure 3a; Scheme S1**). The structure activity relationship showed the free phenol is required for antibiotic activity, while extending the N-alkyl chain preserved its potency. Based on these results we installed an alkyne to facilitate target capture in place of the N-methyl of armeniaspirol, generating probe **2** (**Figure 3b; Scheme S1**). **2** retained activity with an MIC of 8 μg/mL (13.4 μM) against *B. subtilis.*

**Figure 3.**
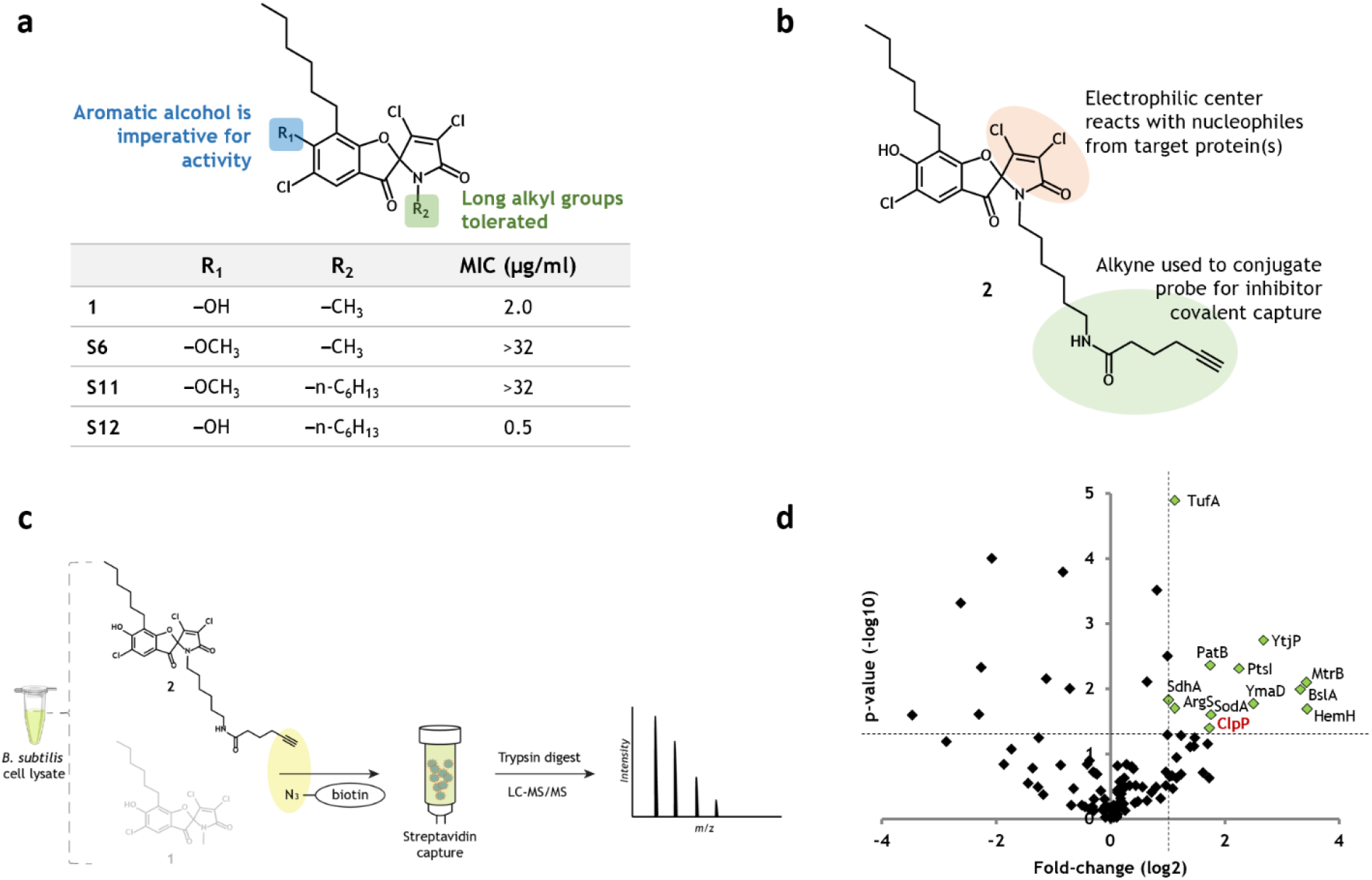
Inhibitor capture strategy for identifying the direct biochemical targets of armeniaspirol. (a) Structure-activity relationship for armeniaspirol antibiotic activity. (b) Armeniaspirol-derived covalent capture probe, **2**. (c) Experimental workflow for profiling armeniaspirol protein targets. The electrophilic center binds to the protein of interest in *B. subtilis* cell lysate. The free alkyne is conjugated to biotin-azide and captured on streptavidin beads. Captured proteins are subjected to digestion and LC-MS/MS for identification. A competition assay with **1** is included to clarify targets. (d) Volcano plot of inhibitor capture, resulting in 12 biologically and statistically significant hits (Log2 fold change > 1, p-value < 0.05).

A competitive chemical proteomics strategy in *B. subtilis* cell lysate was used to identify candidate targets^13^ (**Figure 3c**). Cell lysate was pretreated with **1** to occupy binding sites on potential targets or with a vehicle control to leave binding sites available. Samples were subsequently treated with **2** to capture proteins with accessible binding sites. Captured proteins were recovered by crosslinking the probe with biotin-azide followed by binding onto streptavidin beads. On-bead digestion and LC-MS/MS analysis provided relative abundances of all identified proteins. Twelve proteins showed a statistically significant (p-value <0.05) and biologically relevant (fold-change >2) increase in abundance versus the sample pretreated with the competitor **1** (**Figure 3d**).

Two of the proteins obtained are encoded by essential genes, TufA and ArgS, and both are intimately involved in translation^14–17^. However, our quantitative proteomics and synergy assays suggest translation is not targeted by armeniaspirol. Thus, while armeniaspirol may interact with these proteins, they are not potential targets. Of the remaining proteins, several factors directed us to ClpP, the proteolytic component of the ClpXP protease, as a direct biochemical target. ClpXP plays an integral role in bacterial viability by degrading damaged and misfolded proteins, and is strongly associated with bacterial virulence in multiple Gram-positive pathogens including *S. aureus, S. pneumoniae, and E. faecalis*^18–22^. A further examination of our quantitative proteomics dataset also showed that **1** treatment led to at least a 2-fold enrichment of a number of known or suspected protein substrates of ClpP^23–27^ (**Figure S1**). These data strongly suggest ClpP is an important target of armeniaspirol.

### AAA+ proteases ClpYQ and ClpXP are targets of armeniaspirol

ClpXP and ClpYQ belong to the AAA+ family of proteases in *B. subtilis*^28^. The ClpXP complex is composed of a hexameric ring of the ATPase ClpX bound to a heptameric ring of the serine protease ClpP^29^. ClpXP binds to unstructured peptide tags in proteins, such as the C-terminal tag on the essential cell division protein FtsZ, and targets them for unfolding and proteolysis^30–32^. The ClpYQ complex, also known as HslVU and CodWX, is similarly composed of a hexamer of the ATPase ClpY bound to a hexamer of the protease ClpQ^33–35^. ClpY recognizes and unfolds native protein substrates and passes the unfolded substrate into the catalytic chamber of ClpQ, which performs proteolysis through an N-terminal active site serine^18,33^.

In *B. subtilis* and *S. aureus*, neither ClpP nor ClpQ are encoded by essential genes^14,36^ though in *S. aureus* the double Δ*clpQ* Δ*clpP* mutant is not viable^37^. Furthermore, in the pathogen *Mycobacterium tuberculosis*, which lacks *clpQ, clpP* is essential^38,39^. These observations suggest the combined activity of ClpP and ClpQ are essential. Interestingly, ClpQ was pulled down in one of our chemical proteomic replicates and not detected in the remaining samples. This limited detection is consistent with the low abundance of ClpYQ in *B. subtilis* relative to ClpXP and the low sensitivity of ClpQ to LC-MS-based proteomic detection^40^. We thus hypothesized that armeniaspirol inhibits both the ClpXP and ClpYQ proteases leading to antibiotic activity and sought to test this hypothesis *in vitro* by monitoring both protease and ATP hydrolysis activities (**Figure 4a**).

**Figure 4.**
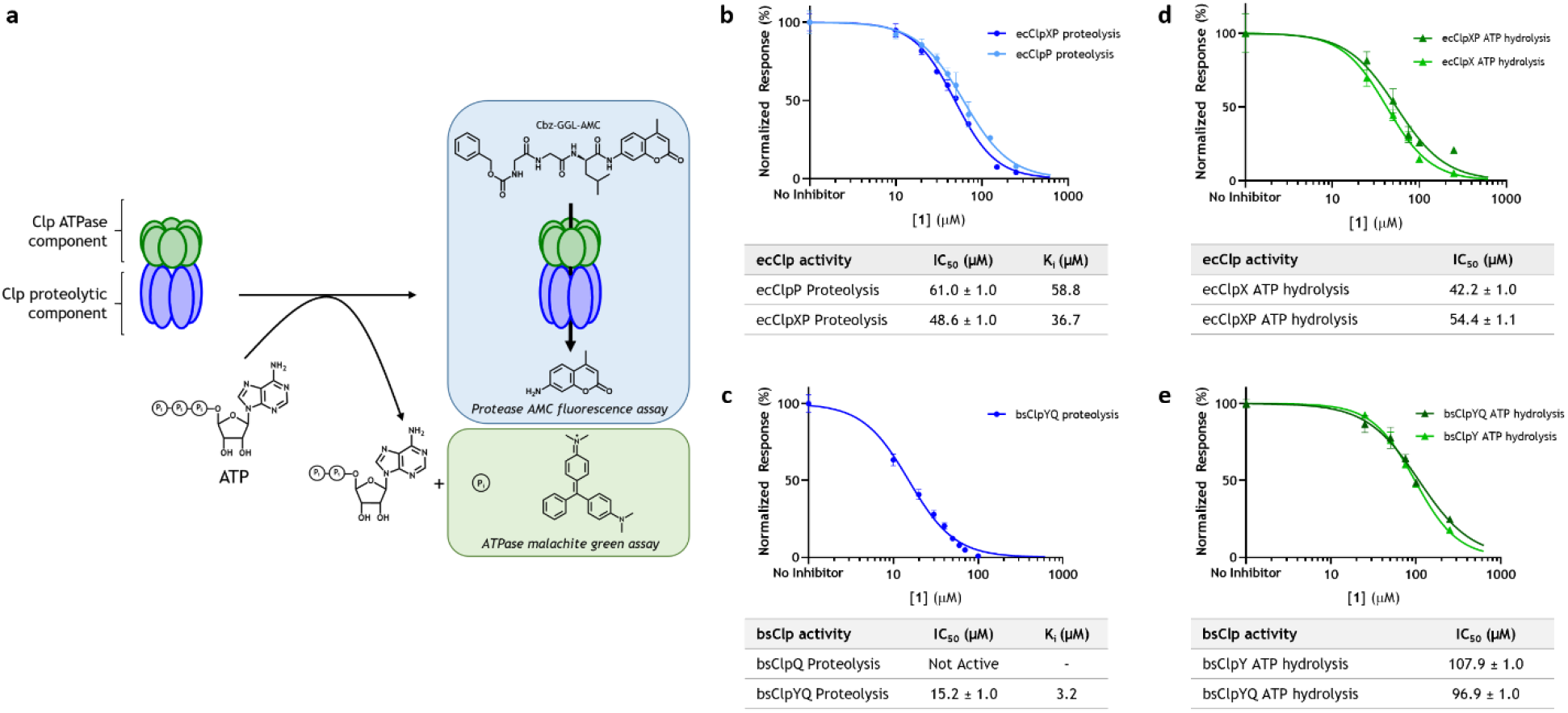
Inhibition of ecClpXP and bsClpYQ by 1. (a) General schematic of the Clp peptidase assay. Degradation of a fluorogenic peptide is observed by the excitation and emission of AMC release. ATP hydrolysis is assayed by malachite green dye-mediated detection of free phosphorous at 650 nm. **1** inhibits proteolysis of (b) ecClpXP and (c) bsClpYQ with an IC_50_ of 48.6 μM with 0.1 mM Suc-LY-AMC and an IC_50_ of 15.2 μM with 0.1 mM Cbz-GGL-AMC, respectively. **1** inhibits ATP hydrolysis of (d) ecClpXP and (e) bsClpYQ with an IC_50_ of 54.4 μM and 96.9 μM, respectively with 1 mM ATP.

Kinetic characterization of recombinant purified ClpXP and ClpYQ showed **1** to be a competitive inhibitor of proteolysis (**Figure S2, Figure S3**). Although we were able to express and purify *B. subtilis* ClpXP (bsClpXP), we could only achieve single turnover for proteolysis under all assay conditions investigated. We thus turned to the homologous *E. coli* ClpXP (ecClpXP), which has been previously characterized *in vitro*^41^. Our results showed **1** inhibited ecClpXP hydrolysis of Suc-LY-AMC *in vitro* with an IC_50_ of 48.6 ± 1.0 μM and a K_i_ of 36.7 μM (**Figure 4b, Figure S2**). In the absence of ClpX, ClpP is known to possess modest peptidase activity and an identical assay showed **1** inhibited ecClpP with an IC_50_ of 61.0 ± 1.0 μM and a K_i_ of 58.8 μM (**Figure 4b, Figure S2**). As an *in vitro* activity assay for *B. subtilis* ClpYQ (bsClpYQ) had previously been developed^33,42^, we readily expressed and purified bsClpQ and bsClpY and were able to kinetically quantify its proteolysis activity with the fluorogenic peptidic substrate, Cbz-GGL-AMC. The addition of **1** showed dose-dependent inhibition of proteolysis with an IC_50_ of 15.2 ± 1.0 μM and a K_i_ of 3.2 μM (**Figure 4c, Figure S3**). Unlike ecClpP, which possesses peptidase activity alone, bsClpQ displayed no activity in the absence of the ATPase component ClpY, and expectedly the complex displayed no activity in the absence of ATP (**Figure S3b**). Interestingly, **1** inhibited ATPase activity of bsClpYQ and ecClpXP with IC_50_s of 96.9 ±

Armeniaspirol however does not appear to inhibit other *B. subtilis* AAA+ proteases, Lon and FtsH. Unlike ClpXP and ClpYQ, these are single polypeptide ATP-dependent proteases that form a hexameric complex to unfold and degrade proteins. LonA is largely responsible for degrading misfolded proteins in *E. coli* by recognition of exposed hydrophobic sequences^43,44^, but plays a minor role in *B. subtilis* protein degradation^45^. In *B. subtilis*, expression of the *fla-che* operon is activated by a known target of LonA, SwrA^46,47^. Reduced proteolysis of SwrA by an inhibited LonA should thus lead to an increase in *fla-che* operon encoded proteins, which is not observed (CheA and CheC decrease by 3.6 and 9.3 fold with p-values of 0.03 and 0.0007, respectively) (**Supplementary File 1**). The other AAA+ protease, FtsH, is a highly selective, membrane bound metalloprotease targeting the cell division protein EzrA and proteins involved in sporulation^48–50^. Our quantitative proteomics shows a significant decrease in EzrA (23 fold decrease, p-value of 0.01) (**Supplementary File 1**). If **1** was inhibiting FtsH, EzrA abundance would be expected to increase.

Individual *B. subtilis* Δ*clpP* and Δ*clpQ* deletion strains (BKK34540 [*clpP::kan*] and BKE16150 [*clpQ::erm*], respectively)^51^ were equally as sensitive to **1** as wild-type *B. subtilis*, determined by MICs of 2 μg/mL. This result is consistent with our hypothesis that antibiotic activity is conferred through a combined mechanism targeting both AAA+ proteases. To validate this, we compared the proteome of wild-type *B. subtilis* treated with **1** to the proteomes of the individual untreated Δ*clpP* and Δ*clpQ* deletion strains. Dimethyl isotopic labelling was used to quantitatively profile changes to the bacterial proteomes, with untreated wild-type *B. subtilis* used as the control (**Figure S4, Supplementary Files 1-3**). Proteins with high false discovery rate confidence with at least 2 or more peptides detected were used in the data analysis. To further filter sparsely quantified proteins, only protein hits that were detected in two of three replicates in either the test or control group were used for the analysis. Missing values were imputed using a missing not at random strategy and the fold change of each mutant relative to wild-type and the statistical significance of this change was determined^52^. Protease substrates are expected to increase in abundance in the protease deletion mutants and **1**-treated samples. Thus we examined the proteins with a two-fold increase in abundance and a p-value ≤ 0.05 (**Figure 5a, Supplementary File 4**). Treatment with **1** led to an increase in 109 proteins. The *ΔclpP* and *ΔclpQ* mutants showed 264 and 191 proteins increased, respectively. The 41 proteins that overlap between **1** treatment and the *ΔclpP* mutant and the 21 proteins that overlap between **1** treatment and the *ΔclpQ* mutant are a statistically significant increase over the expected values from random as determined by the Chi square goodness of fit test (P < 0.0001). This strongly supports our hypothesis that armeniaspirol is inhibiting the ClpXP and ClpYQ proteases *in vivo* in *B. subtilis*.

**Figure 5.**
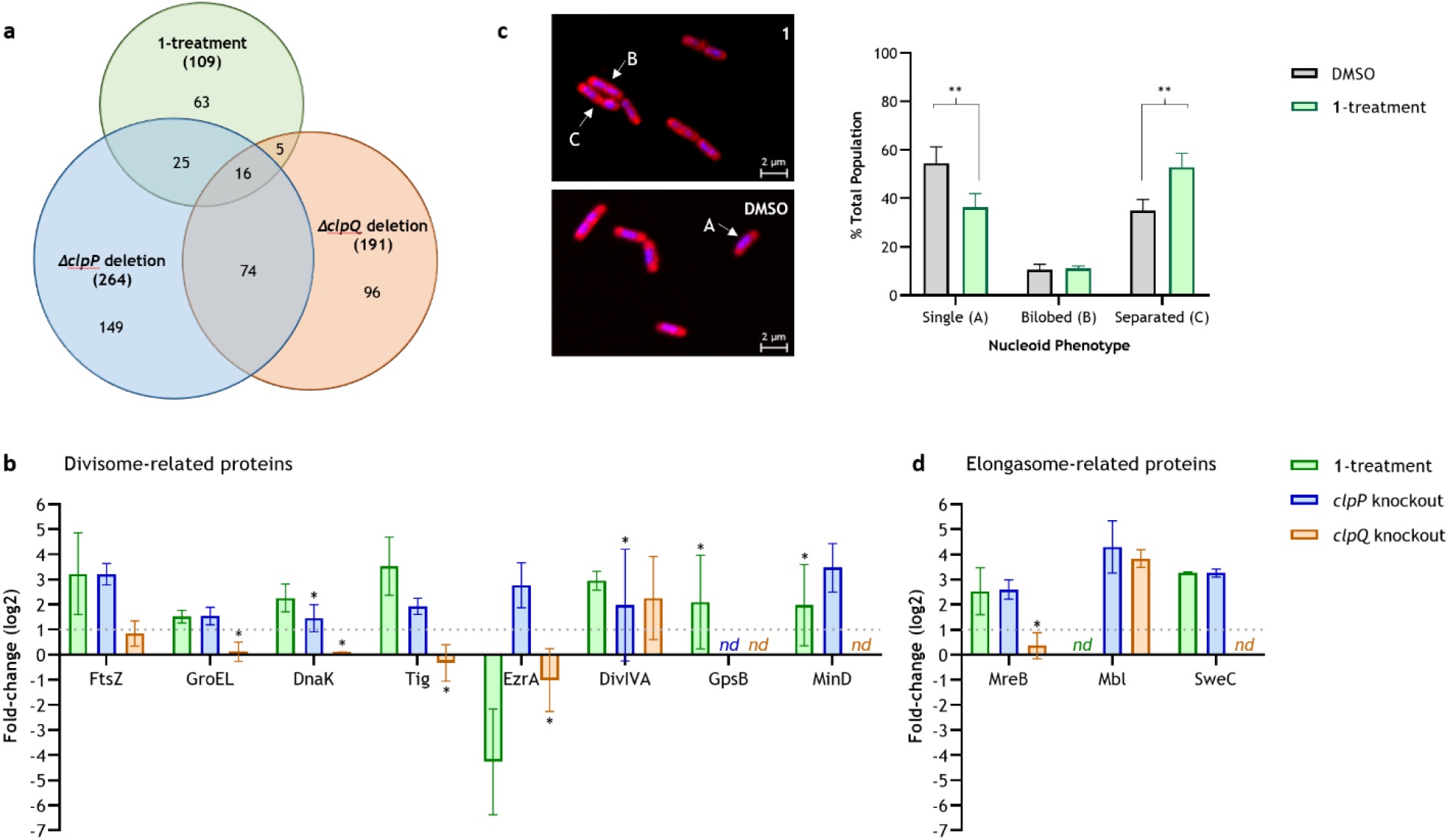
Dysregulation of the divisome and elongasome complexes in *B. subtilis* by 1-treatment and Δ*clpP* and Δ*clpQ* mutants. (a) Venn diagram depicting upregulated proteins across treatments obtained from biological triplicates (Log2 fold change > 1, p-value < 0.05). (b) Effects on divisome and elongasome protein abundances according to treatment (*nd;* not detected). *not significant, detected in 2 out of 3 replicates. (c) Micrographs of **1**-treated versus vehicle control treated *B. subtilis* showing representations of single nucleoids (A), bilobed nucleoids (B), and separated nucleoids (C). Septation defects were graphed according to nucleoid phenotype (biological triplicate; DMSO n=149, **1**-treatment n=157). **p < 0.05, statistical analysis was performed by 2-way ANOVA with Holm-Sidak's multiple comparison test.

### Armeniaspirol dysregulates the B. subtilis divisome

Examination of the enriched proteins after treatment with **1** was used to identify the mechanism by which inhibition of ClpXP and ClpYQ leads to antibiotic activity. Gene ontology (GO) analysis of the enriched proteins identified indirect effects of treatment with a bactericidal antibiotic^53,54^. For example, the enriched proteins showed an overrepresentation of enzymes involved in branched amino acid biosynthesis, glycolytic and coenzyme metabolic processes, and nucleotide scavenging metabolism consistent with bactericidal antibiotic treatment. However, no clear direct effect leading to antibiotic activity was identified from the GO analysis.

Manual examination of each enriched protein showed several cell division related components had increased in abundance upon treatment with **1** (**Figure 5b**). The key divisome protein FtsZ increased significantly upon treatment with **1**. FtsZ polymerizes at the mid-cell forming a dynamic ring upon which additional divisome proteins are scaffolded to guide septum synthesis between daughter cells^55,56^. Additionally the highly abundant chaperones GroEL, DnaK, and Tig, known to play a role in FtsZ function, also increased^40,57–62^. As these are known ClpXP substrates^23–25,27,57^, they show a significant increase in the *ΔclpP* dataset as well (**Figure 5b, Figure S1**).

FtsZ assembles with FtsA, ZapA, and EzrA into the early divisome, which recruits DivIVA and a number of extracytoplasmic proteins to form the late divisome^63^. Our quantitative proteomics shows a significant increase in the negative regulator of cell division, DivIVA, as well as an increase in two of the three replicates of its ortholog GpsB^64^ upon treatment with **1** (**Figure 5b**). DivIVA is recruited to sites of negative membrane curvature, like the polar regions and nascent division sites, where it can complex with MinC and MinD to inhibit FtsZ polymerization^65–69^. Furthermore, DivIVA is known to inhibit cell division during competency^70^. Independent of the Min system, DivIVA appears to play a key role in coordinating chromosome segregation and cell division presumably in conjunction with the SMC protein^59,71^.

As DivIVA is constitutively expressed with low transcript levels, its significant increase in abundance upon treatment with **1** is post-transcriptional and likely due to decreased proteolysis^72^. The quantitative proteomic dataset from the Δ*clpQ* mutant also shows a large significant increase in DivIVA suggesting it may be a direct target of the ClpYQ protease. Intriguingly DivIVA is proposed to be a substrate of ClpXP in *Streptococcus mutans*, which lacks *clpQ^73^.* It is thus possible that ClpXP plays a minor role in degrading DivIVA in *B. subtilis*, supported by the increase in DivIVA abundance in two out of three replicates of the *ΔclpP* mutant dataset. Few targets of the ClpYQ protease are known in *B. subtilis* and none are involved in cell division, proposing a significant new role for ClpYQ^74^.

The tight regulation of the divisome Z-ring formation is further maintained by key negative regulators of cell division, including the Min system comprised of MinC and MinD, which are involved in septum placement at the midcell^75^. MinD increases in abundance with treatment by **1**, though it is only detected in two of the three replicates. However, MinD increases significantly in the *ΔclpP* dataset (**Figure 5b, Table S3**). Like DivIVA, the *min* operon is constitutively expressed and not differentially regulated as the cell progresses through the cell cycle^72^. Thus, the observed significant increase in MinD in the *ΔclpP* dataset and the increase seen in two replicates from treatment with **1** are expected to be due to inhibition of ClpXP. While MinD is known to be a client of ClpXP in *E. coli*, it has not previously been characterized as a *B. subtilis* ClpXP client^23,25^. However, our work clearly supports this hypothesis.

Dysregulation of the divisome can be catastrophic to cell division. For example, overexpression of FtsZ without FtsA in *E. coli* is known to block cell division^76–79^, overexpression of DivIVA in *B. subtilis* appears to be not viable^80^, and a 15-fold increase in MinCD levels or overexpression of a MinD-GFP fusion inhibits Z-ring formation and leads to cell death^68,81^. As treatment with **1** lead to increases in all of these key divisome components, we hypothesized that it should result in division defects. We examined the impact of **1** treatment on cell division. Confocal microscopy of DNA (DAPI) and membrane (FM 4-64FX) stained wild-type *B. subtilis* treated with a sub-lethal dose of **1** (½ MIC) showed a statistically significant increase in the number of two nucleoid containing cells and a decrease in single nucleoid containing cells compared to untreated cells (**Figure 5c; Figure S5**). These observations are in accordance with a defect in cell division, and consistent with our proposed mechanism involving a negative regulation of cell division by blocking ClpYQ and ClpXP protease activities.

In addition to the divisome, rod shaped bacteria like *B. subtilis* contain a related structure called the elongasome, which is responsible for peptidoglycan hydrolysis and synthesis along the cell axis allowing for cylindrical growth^82^. The elongasome is built upon of polymer of MreB, or its homologs Mbl and MreBH, which tightly associate to the membrane^83–85^. This polymer anchors a complex of integral membrane proteins, such as FtsE and FtsX^86,87^, and extracellular proteins, such as CwlO^88^, a peptidoglycan hydrolase, and multiple PBPs^89,90^, which synthesize new peptidoglycan. Armeniaspirol also appears to dysregulate the elongasome since treatment with **1** leads to a significant increase in MreB (Mbl and MreBH are not detected). As MreB is constitutively expressed^72^, its increase is presumably due to inhibition of proteolysis. Consistent with this, the *ΔclpP* mutant shows a significant increase in MreB and Mbl (MreBH is not detected) and the *ΔclpQ* mutant shows a significant increase in Mbl (**Figure 5d, Table S3**), suggesting that these key elongasome components are substrates of the Clp proteolytic machinery. MreB is a known inhibitor of cell division in many bacteria, and over-expression of MreB and MreBH is lethal in *B. subtilis*^90–93^.

One of the proteins that increases in abundance the most with **1** treatment is SweC (also called YqzC)^6^. SweC and its binding partner SweD interact with the FtsE-FtsX integral membrane protein complex of the elongasome and acts as a mechanotransmission domain to activate the extracellular D,L-endopeptidase CwlO, which then degrades the distal load bearing peptidoglycan layers^88,94^. Overactivation of peptidoglycan hydrolases, such as CwlO, can lead to weakening of the peptidoglycan and cell lysis^95^. While *sweC* expression is repressed by Spo0A under sporulation conditions, proteins whose transcription is comparably repressed by Spo0A, such as PurT^96^, show no significant increase with **1** treatment. This suggests that the observed increase in SweC abundance may be due to inhibition of its proteolysis. Consistent with that hypothesis, the Δ*clpP* mutant shows a significant increase in SweC. Thus, in addition to arresting cell division though dysregulation of the divisome, armeniaspirol may dysregulate the elongasome and lead to over-activation of peptidoglycan hydrolases, weakening the cell wall.

## Discussion

The Gram-positive antibiotic armeniaspirol inhibits both the ClpXP and ClpYQ AAA+ proteases, enriching client proteins of these proteases that consequently disrupt cell division. This new mechanism of action is rooted in analysis of two key proteomic data sets. Our chemical proteomic data set implicates ClpXP and ClpYQ as direct biochemical targets of armeniaspirol, and our quantitative proteomics data sets demonstrate an impact on the divisome. It is therefore essential to critically and thoroughly evaluate these analyses.

The competition-dependent chemical proteomics assay suggests that ClpP is a direct biochemical target of armeniaspirol. ClpP shows a 3.3-fold increase in abundance in the inhibitor capture experiment. HemH, MtrB, and BslA all showed 10-fold or greater enrichment. All three are encoded by non-essential genes and no additional data or precedent supports them as potential direct targets of armeniaspirol. HemH is a ferrochelatase, which catalyzes the last step in *B. subtilis* heme biosynthesis^97^. The protein encoded by its mammalian ortholog has been frequently captured as an off-target binder of small molecule kinase inhibitors in chemical proteomic experiments^98^. It is therefore proposed to have a promiscuous protophorphorin binding site, which likely binds the probe in our capture experiments. Thus, while HemH may bind armeniaspirol non-selectively, there is no clear mechanism connecting HemH inhibition and antibiotic activity. In fact, *S. aureus hemH* mutants are selected for under gentamicin treatment, confirming their viability^99^. MtrB, a tryptophan operon RNA-binding attenuation protein (TRAP), regulates *B. subtilis* expression of the tryptophan biosynthesis operon. MtrB allosterically binds Trp leading to premature transcriptional termination of *trpEDCFBA* operon^100^. Inhibition of MtrB by RtpA, an anti-TRAP protein, leads to high levels of TrpE expression and activity^101^. Our quantitative proteomics analysis of armeniaspirol treatment does not show a statistically significant change in abundance of TrpE, suggesting armeniaspirol does not inhibit MtrB. Lastly, BslA is an extracellular self-assembling hydrophobin protein that coats *B. subtilis* biofilms^102^. It has an extremely hydrophobic cap region that presumably can interact with the probe enabling non-covalent capture and was thus eliminated as a direct target.

Another five proteins from the chemical proteomics experiment were enriched at comparable levels to ClpP: YtjP (6.4-fold), YmaD (5.7-fold), PtsI (4.7-fold), SodA (3.4-fold), and PatB (3.4-fold). These proteins could not be unambiguously eliminated as potential direct biochemical targets, though they are all encoded by non-essential genes. Both PtsI, which is Enzyme 1 from the phosphotransferase system, and SodA, superoxide dismutase, are two of the most abundant proteins in the *B. subtilis* proteome^40^. Their high concentrations may facilitate pull down via non-selective interactions with the probe. YtjP is a metalloprotease of which very little is known. Based on homology to the *S. aureus* ortholog, the holo form of YtjP likely contains an exposed cysteine that that may react with the electrophilic probe for capture. Similarly YmaD is a peroxiredoxin related to *E. coli* OsmC that has a hydrophobic binding cavity containing a highly nucleophilic cysteine, potentially enabling capture by the probe^103^. Lastly, PatB, a cystathione-β-lyase (CBL) responsible for methionine biosynthesis, has overlapping function with MetC and a double mutant of both is unable to grow on cystathione as the sole sulfur-source^104^. CBL from a variety of bacteria are inactivated by cysteine alkylation with N-ethylmaleimide^105^, thus it is possible that the probe is covalently modifying PatB. However, as inhibition of CBL does not appear to correlate with antimicrobial activity^106^, PatB was downgraded to an unlikely candidate for the direct biochemical target of armeniaspirol. Thus, while we cannot unambiguously rule out these five enriched proteins, only ClpP, which is known to be essential when its homolog ClpQ is not present, is well supported as a potential direct biochemical target. Our subsequent *in vitro* kinetic characterization clearly demonstrates that both the ClpXP and ClpYQ complexes are directly and competitively inhibited by armeniaspirol, and that treatment of *B. subtilis* leads to an increase in abundance of proteins known to be ClpXP substrates.

Particularly unexpected from our kinetic characterization was the observation that armeniaspirol *competitively* inhibits ClpXP and ClpYQ in biochemical assays. As these targets were identified through a covalent inhibitor capture chemical proteomics strategy, irreversible inhibition would have been expected^13^. Two hypotheses are consistent with covalent target capture with competitive biochemical inhibition. If armeniaspirol is a reversible covalent inhibitor, it is possible to both capture the target and to exhibit competitive inhibition kinetics^107^. However the Michael addition/elimination-based substitution chemistry that armeniaspirol and related β-chloro-α,β-unsaturated lactams undergo is inconsistent with reversible protein alkylation^5,108^. Alternatively, a very slow inactivation rate (*k*_inact_) following formation of the non-covalent enzyme-inhibitor complex can enable covalent capture but be insufficient to covalently modify enough enzyme in biochemical assays to be able to exhibit detectable irreversible inhibition. A clear prediction from this hypothesis is a very low level of target capture in the chemical proteomics. This is fully consistent with our data, where very few proteins are captured, with only twelve biologically and statistically significant proteins being identified in the capture experiment (**Figure 3d**). Furthermore, based on the capture of the natively biotinylated protein AccB, which is known to have a similar cytosolic abundance to ClpP, it appears that significantly less than 1% of ClpP is captured, consistent with a very slow *k*_inact_^40^. We thus propose that while armeniaspirol is electrophilic, its ability to alkylate proteins does not play a significant role in its inhibition of ClpXP and ClpYQ. Intriguingly based on this analysis, it may be possible to identify armeniaspirol inspired compounds that retain potent ClpXP/ClpYQ inhibition without the liability of an electrophilic center.

Critical analysis of our quantitative proteomics data suggests armeniaspirol perturbs the divisome and elongasome through enrichment of key proteins FtsZ, DivIVA, and MreB. Increased abundance in each of these is known to inhibit cell division^76–80,90–93^. No other clear link to antibiotic activity could be identified from the quantitative proteomics data set. Of the 109 proteins that are shown to significantly increase in abundance with armeniaspirol treatment, 24 are from the group of 100 most abundant proteins in *B. subtilis*^57^. This is a highly significant enrichment of the most abundant proteins (p < 0.00001, Chi square goodness of fit test). Inhibition of ClpXP and ClpYQ by armeniaspirol provides a potential mechanism for this enrichment by reduction of protein quality control via proteolysis.

Seventeen translation related proteins increase in abundance upon treatment with armeniaspirol. For example, ribosomal proteins required for the assembly of the large subunit, L12 and L13, and small subunit, S4, S11, and S13 are increased, suggesting assembly may be perturbed^109^. The ribosome checkpoint GTPases Obg and RgbA are also significantly increased in abundance, and in the presence of elevated (p)ppGpp from the stringent response, these are known to inhibit ribosome assembly^110^. We propose this is a general, indirect effect of inhibition of proteolysis by armeniaspirol. The stringent response, which is initiated by an increase in uncharged tRNA^111^, presumably due to the decrease in proteolysis, is well known to inhibit protein synthesis until sufficient charged tRNAs are available^112^. Supporting a stringent response is the increase in abundance of several aminoacyl-tRNA synthetases, and the observation that many of the most abundant *B. subtilis* proteins show an increased abundance upon treatment with armeniaspirol, reducing the available amino acid pool. Furthermore, if armeniaspirol was directly inhibiting ribosome assembly and function as its primary mechanism of action, it should be synergistic with the protein synthesis inhibitor tetracycline. However, our data shows that armeniaspirol and tetracycline are indifferent (FIC = 0.625; **Table S2**). Thus, while several factors suggest a perturbation of translation by armeniaspirol, this is likely an indirect effect of inhibition of ClpXP and ClpYQ via the stringent response.

Our quantitative proteomics also show six iron transport proteins significantly increase in abundance in the armeniaspirol treatment. The elemental iron transporter EfeO, the Fe-citrate transporter YfmC, and the siderophore transporters YclQ, FhuD, YfiY, FeuA are all increased. While upregulation of the genes encoding these proteins is typical of iron limitation^113^, there is a high coincidence of gene regulation between iron limitation and the stringent response^114^, suggesting that this may also be an indirect effect of armeniaspirol treatment via the stringent response. However, we cannot rule out this effect being mediated by armeniaspirol, perhaps via binding to HemH. Regardless, there is little support for this perturbation of iron homeostasis, especially in iron-sufficient media, as a mechanism of action.

Thus critical analysis of our quantitative proteomics data supports armeniaspirol acting though perturbation of the divisome and elongasome. Consistent with this mechanism we observe a defect in cell division upon treatment with armeniaspirol. The quantitative proteomics data suggests armeniaspirol treatment initiates a stringent response, presumably via reduction in the pool of amino acylated-tRNAs due to reduced proteolysis. This is supported by the observation that many of the most abundant *B. subtilis* proteins increase in abundance upon treatment with armeniaspirol, reducing the available amino acid pool.

Our analysis thus shows that disrupting the divisome and elongasome via inhibition of AAA+ proteases function is a new strategy for antibiotic activity. Many antibiotic discovery efforts have focused on the divisome because of its essential nature in bacterial cell division. An obvious target has been to develop inhibitors of FtsZ, because inhibition of this highly conserved essential protein should lead to broad-spectrum antibiotic activity^115^. While potent *in vitro* and *in vivo* inhibitors have been discovered, spontaneous mutations of FtsZ leading to resistance is a serious and limiting concern^116^. Thus targeting of AAA+ proteases by armeniaspirol leading to disruption of divisome function is particularly appealing since laboratory selection efforts have failed to generate armeniaspirol-resistant strains^4^. In addition to disrupting the divisome, armeniaspirol dysregulates the elongasome as well. There are strong parallels between the divisome and elongasome^82^. Both complexes coordinate lipid II synthesis with peptidoglycan hydrolysis and synthesis through scaffolding of integral membrane proteins and extracellular proteins on a cytosolic membrane bound polymer^117,118^. In fact, the divisome and elongasome are thought to be evolutionarily related complexes. It is thus particularly interesting that ClpXP and ClpYQ appear to play an overlapping role in regulating the function of each via proteolysis of, at minimum, key proteins FtsZ from the divisome and MreB from the elongasome.

Our work identifies armeniaspirol as the first natural product inhibitor of the AAA+ protease ClpXP. A number of natural products are known to target the AAA+ proteases. These include the 20S proteasomes inhibitors such as epoxomicin and related α,β-epoxyketones as well as salinosporamide and similar β-lactone proteasome inhibitors^119–121^. Both of these families of compounds covalently modify the N-terminal catalytic Thr of the 20S proteasome in actinobacteria and eukaryotes^122^ but have not been shown to inhibit ClpP or ClpQ. The acyldepsipeptide (ADEP) natural product antibiotics bind reversibly to ClpP, activating the protease even in the absence of ClpX^123^. This over-activation of ClpP leads to uncontrolled protein degradation and antibiotic activity^124^. Cyclomarin A, lassomycin, and ecumicin all appear to bind the N-terminal domain of the ClpC ATPase^123^. Cyclomarine A leads to uncontrolled proteolysis whereas lassomycin and ecumicin increase ClpCP ATP hydrolysis but inhibits proteolysis^125–127^. Armeniaspirol is distinct from all of these, binding both ClpXP and ClpYQ and competitively inhibiting both ATP hydrolysis and proteolysis of the complexes and the individual catalytic domains.

ClpXP and ClpYQ have fundamentally different proteolytic machinery. ClpP is a serine protease with Ser-His-Asp catalytic triad^128,129^. ClpQ is a homolog of the β-subunits of the eukaryotic, archaebacterial, and actinobacterial 20S proteasomes and possesses an N-terminal catalytic Ser^130^. The ATPases ClpY and ClpX however are very similar. Both are members of the Clp/HSP100 family^131,132^. Based on the homology of the ATPases, a reasonable hypothesis for armeniaspirol's inhibition of both complexes would be binding to the conserved ATPase domains, inhibiting function of the complex. However, armeniaspirol's ability to inhibit ClpP function alone and the capture of ClpP in our chemical proteomics suggests armeniaspirol may have a more complex interaction with the AAA+ proteases, potentially binding at the interface between protease and ATPase. Characterizing the molecular mechanism of binding to these complexes will thus be of particular interest as it will identify a key new site for engaging these pharmacologically relevant targets.

ClpXP has been identified as a promising target for drug development due to its significant role in regulating virulence in diverse pathogens^19–22,133,134^. Pioneering work has uncovered synthetic β-lactone-, phenyl ester-, and boronate-based inhibitors of ClpXP^135–137^. While promising, significant challenges including resistance, stability, and potential reduction of susceptibility to other antibiotics exist for these ClpP inhibitors^136,138–140^. Our discovery of armeniaspirol as a dual inhibitor of ClpXP and ClpYQ is thus particularly noteworthy. While armeniaspirol can attenuate virulence via ClpXP inhibition, it additionally targets ClpYQ conferring antibiotic activity unlike current ClpP inhibitors.

Lastly, under laboratory selection experiments, it has not been possible to raise armeniaspirol resistant strains^4^. This is significantly different from other natural products that perturb AAA+ protease activity, such as the ADEPs, which rapidly elicit resistant phenotypes^141,142^. While engaging multiple targets typically reduces the probability of resistance arising^143^, this is only true if both targets have independent essential function. As the armeniaspirol mechanism requires inhibition of both ClpXP and ClpYQ, resistance should arise since a mutation of either target that decreases armeniaspirol inhibition will lead to resistance. Clearly the undetectably low frequency of resistance suggests additional mechanisms are at work with this antibiotic, as is typical of natural products antibiotics^144^. It thus remains an exciting ongoing question as to what additional targets armeniaspirol interacts with, preventing the emergence of resistant strains.

In summary we have demonstrated that armeniaspirol inhibits the AAA+ proteases ClpXP and ClpYQ. This work identifies armeniaspirol as the first natural product inhibitor of the coveted antivirulence target ClpP. Unlike known inhibitors and activators of ClpP, armeniaspirol also inhibits ClpYQ, and this dual inhibition of ClpXP and ClpYQ leads to antibiotic activity. Furthermore, it has not been possible to select for armeniaspirol resistant strains. We show armeniaspirol competitively inhibits ClpXP and ClpYQ, suggesting that while it possesses an undesirable electrophilic center, this reactivity is not necessary for its function and may be eliminated through medicinal chemistry efforts. In addition, our work shows that inhibition of ClpXP and ClpYQ dysregulates key proteins of the divisome and elongasome by preventing their targeted proteolysis, leading to an arrest in cell division and we provide preliminary data identifying a number of new targets for these proteases, including DivIVA and Mbl as potential targets of the uncharacterized ClpYQ protease. These results clearly elevate armeniaspirol and its new mechanism of action to a highly promising position for antibiotic development.

## Acknowledgements

*B. subtilis* knockout strain MGNA-A086 was ordered from National BioResource Project (NIG, Japan): *B. subtilis. B. subtilis* knockout strains BKE16150 and BKK34540 were ordered from the Bacillus Genome Stock Centre (BGSC). MRSA strains were provided by Dr. Wilmara Salgado-Pabón (University of Iowa). Expression plasmids harbouring *E. coli clpX* and *clpP* genes were generously provided by Dr. Walid Houry (University of Toronto). We thank Dr. Zoran Minic of the John L. Holmes Mass Spectrometry Facility (University of Ottawa) for proteomics support.

## References

(1) Brown, E. D.; Wright, G. D. Antibacterial drug discovery in the resistance era. Nature 2016, 529 (7586), 336–343.

(2) The 10 x ' 20 initiative: pursuing a global commitment to develop 10 new antibacterial drugs by 2020. Clin. Infect. Dis. 2010, 50 (8), 1081–1083.

(3) Talbot, G. H.; Jezek, A.; Murray, B. E.; Jones, R. N.; Ebright, R. H.; Nau, G. J.; Rodvold, K. A.; Newland, J. G.; Boucher, H. W. The Infectious Diseases Society of America's 10 × '20 initiative (10 new systemic antibacterial agents US Food and Drug Administration approved by 2020): is 20 × '20 a possibility? Clin. Infect. Dis. 2019, 69 (1), 1–11.

(4) Dufour, C.; Wink, J.; Kurz, M.; Kogler, H.; Olivan, H.; Sablé, S.; Heyse, W.; Gerlitz, M.; Toti, L.; Nußer, A.; et al. Isolation and structural elucidation of armeniaspirols A-C: Potent antibiotics against gram-positive pathogens. Chem. - A Eur. J. 2012, 18 (50), 16123–16128.

(5) Couturier, C.; Bauer, A.; Rey, A.; Schroif-Dufour, C.; Broenstrup, M. Armeniaspiroles, a new class of antibacterials: antibacterial activities and total synthesis of 5-chloro-Armeniaspirole A. Bioorg. Med. Chem. Lett. 2012, 22 (19), 6292–6296.

(6) Michna, R. H.; Commichau, F. M.; Tödter, D.; Zschiedrich, C. P.; Stülke, J. SubtiWiki-A database for the model organism Bacillus subtilis that links pathway, interaction and expression information. Nucleic Acids Res. 2014, 42 (D1), 692–698.

(7) Boersema, P. J.; Raijmakers, R.; Lemeer, S.; Mohammed, S.; Heck, A. J. R. Multiplex peptide stable isotope dimethyl labeling for quantitative proteomics. Nat. Protoc. 2009, 4 (4), 484–494.

(8) Stepanek, J. J.; Lukežič, T.; Teichert, I.; Petković, H.; Bandow, J. E. Dual mechanism of action of the atypical tetracycline chelocardin. Biochim. Biophys. Acta - Proteins Proteomics 2016, 1864 (6), 645–654.

(9) Wenzel, M.; Patra, M.; Albrecht, D.; Chen, D. Y. K.; Nicolaou, K. C.; Metzler-Nolte, N.; Bandow, J. E. Proteomic signature of fatty acid biosynthesis inhibition available for in vivo mechanism-of-action studies. Antimicrob. Agents Chemother. 2011, 55 (6), 2590–2596.

(10) Wenzel, M.; Kohl, B.; Münch, D.; Raatschen, N.; Albada, H. B.; Hamoen, L.; Metzler-Nolte, N.; Sahl, H.-G.; Bandow, J. E. Proteomic Response of Bacillus subtilis to Lantibiotics Reflects Differences in Interaction with the Cytoplasmic Membrane. Antimicrob. Agents Chemother. 2012, 56 (11), 5749–5757.

(11) Bandow, J. E.; Brötz, H.; Leichert, L. I. O.; Labischinski, H.; Hecker, M. Proteomic approach to understanding antibiotic action. Antimicrob. Agents Chemother. 2003, 47 (3), 948–955.

(12) Sender, U.; Bandow, J.; Engelmann, S.; Lindequist, U.; Hecker, M. Proteomic signatures for daunomycin and adriamycin in Bacillus subtilis. Pharmazie 2004, 59 (1), 65–70.

(13) Wright, M. H.; Sieber, S. A. Chemical proteomics approaches for identifying the cellular targets of natural products. Nat. Prod. Rep. 2016, 33 (5), 681–708.

(14) Peters, J. M.; Colavin, A.; Shi, H.; Czarny, T. L.; Larson, M. H.; Wong, S.; Hawkins, J. S.; Lu, C. H. S.; Koo, B.-M.; Marta, E.; et al. A comprehensive, CRISPR-based functional analysis of essential genes in bacteria. Cell 2016, 165 (6), 1493–1506.

(15) Sprinzl, M. Elongation factor Tu: a regulatory GTPase with an integrated effector. Trends Biochem. Sci. 1994, 19 (6), 245–250.

(16) Harvey, K. L.; Jarocki, V. M.; Charles, I. G.; Djordjevic, S. P. The diverse functional roles of elongation factor tu (Ef-tu) in microbial pathogenesis. Front. Microbiol. 2019, 10, 1–19.

(17) Giegé, R.; Springer, M. Aminoacyl-tRNA synthetases in the bacterial world. EcoSal Plus 2016, 7 (1), 1–83.

(18) Olivares, A. O.; Baker, T. A.; Sauer, R. T. Mechanistic insights into bacterial AAA+ proteases and protein-remodelling machines. Nat. Rev. Microbiol. 2016, 14 (1), 33–44.

(19) Mei, J.; Nourbakhsh, F.; Ford, C. W.; Holden, D. W. Identification of Staphylococcus aureus virulence genes in a murine model of bacteraemia using signature-tagged mutagenesis. Mol. Microbiol. 1997, 26 (2), 399–407.

(20) Frees, D.; Qazi, S. N. A.; Hill, P. J.; Ingmer, H. Alternative roles of ClpX and ClpP in Staphylococcus aureus stress tolerance and virulence. Mol. Microbiol. 2003, 48 (6), 1565–1578.

(21) Kwon, H. Y.; Ogunniyi, A. D.; Choi, M. H.; Pyo, S. N.; Rhee, D. K.; Paton, J. C. The ClpP protease of Streptococcus pneumoniae modulates virulence gene expression and protects against fatal pneumococcal challenge. Infect. Immun. 2004, 72 (10), 5646–5653.

(22) Cassenego, A. P. V.; de Oliveira, N. E. M.; Laport, M. S.; Abranches, J.; Lemos, J. A.; Giambiagi-deMarval, M. The CtsR regulator controls the expression of clpC, clpE and clpP and is required for the virulence of Enterococcus faecalis in an invertebrate model. Antonie van Leeuwenhoek, Int. J. Gen. Mol. Microbiol. 2016, 109 (9), 1253–1259.

(23) Flynn, J. M.; Neher, S. B.; Kim, Y. I.; Sauer, R. T.; Baker, T. A. Proteomic discovery of cellular substrates of the ClpXP protease reveals five classes of ClpX-recognition signals. Mol. Cell 2003, 11 (3), 671–683.

(24) Feng, J.; Michalik, S.; Varming, A. N.; Andersen, J. H.; Albrecht, D.; Jelsbak, L.; Krieger, S.; Ohlsen, K.; Hecker, M.; Gerth, U.; et al. Trapping and proteomic identification of cellular substrates of the ClpP protease in staphylococcus aureus. J. Proteome Res. 2013, 12 (2), 547–558.

(25) Neher, S. B.; Villén, J.; Oakes, E. C.; Bakalarski, C. E.; Sauer, R. T.; Gygi, S. P.; Baker, T. A. Proteomic profiling of ClpXP substrates after DNA damage reveals extensive instability within SOS regulon. Mol. Cell 2006, 22 (2), 193–204.

(26) Miethke, M.; Hecker, M.; Gerth, U. Involvement of Bacillus subtilis ClpE in CtsR degradation and protein quality control. J. Bacteriol. 2006, 188 (13), 4610–4619.

(27) Gerth, U.; Kock, H.; Kusters, I.; Michalik, S.; Switzer, R. L.; Hecker, M. Clp-dependent proteolysis down-regulates central metabolic pathways in glucose-starved Bacillus subtilis. J. Bacteriol. 2008, 190 (1), 321–331.

(28) Sauer, R. T.; Baker, T. A. AAA+ proteases: ATP-fueled machines of protein destruction. Annu. Rev. Biochem. 2011, 80, 587–612.

(29) Baker, T. A.; Sauer, R. T. ClpXP, an ATP-powered unfolding and protein-degradation machine. Biochim. Biophys. Acta - Mol. Cell Res. 2012, 1823 (1), 15–28.

(30) Camberg, J. L.; Hoskins, J. R.; Wickner, S. ClpXP protease degrades the cytoskeletal protein, FtsZ, and modulates FtsZ polymer dynamics. Proc. Natl. Acad. Sci. 2009, 106 (26), 10614–10619.

(31) Camberg, J. L.; Hoskins, J. R.; Wickner, S. The Interplay of ClpXP with the Cell Division Machinery in Escherichia coli. J. Bacteriol. 2011, 193 (8), 1911–1918.

(32) Camberg, J. L.; Viola, M. G.; Rea, L.; Hoskins, J. R.; Wickner, S. Location of Dual Sites in E. coli FtsZ Important for Degradation by ClpXP; One at the C-Terminus and One in the Disordered Linker. PLoS One 2014, 9 (4), e94964.

(33) Kang, M. S.; Lim, B. K.; Seong, I. S.; Seol, J. H.; Tanahashi, N.; Tanaka, K.; Chung, C. H. The ATP-dependent CodWX (HslVU) protease in Bacillus subtilis is an N-terminal serine protease. EMBO J. 2001, 20 (4), 734–42.

(34) Kang, M. S.; Kim, S. R.; Kwack, P.; Lim, B. K.; Ahn, S. W.; Rho, Y. M.; Seong, I. S.; Park, S. C.; Eom, S. H.; Cheong, G. W.; et al. Molecular architecture of the ATP-dependent CodWX protease having an N-terminal serine active site. EMBO J. 2003, 22 (12), 2893–2902.

(35) Rho, S.-H.; Park, H. H.; Kang, G. B.; Im, Y. J.; Kang, M. S.; Lim, B. K.; Seong, I. S.; Seol, J.; Chung, C. H.; Wang, J.; et al. Crystal structure of Bacillus subtilis CodW, a noncanonical HslV-like peptidase with an impaired catalytic apparatus. Proteins 2008, 71 (2), 1020–1026.

(36) Chaudhuri, R. R.; Allen, A. G.; Owen, P. J.; Shalom, G.; Stone, K.; Harrison, M.; Burgis, T. A.; Lockyer, M.; Garcia-Lara, J.; Foster, S. J.; et al. Comprehensive identification of essential Staphylococcus aureus genes using Transposon-Mediated Differential Hybridisation (TMDH). BMC Genomics 2009, 10, 291.

(37) Frees, D.; Thomsen, L. E.; Ingmer, H. Staphylococcus aureus ClpYQ plays a minor role in stress survival. Arch. Microbiol. 2005, 183 (4), 286–291.

(38) Raju, R. M.; Unnikrishnan, M.; Rubin, D. H. F.; Krishnamoorthy, V.; Kandror, O.; Akopian, T. N.; Goldberg, A. L.; Rubin, E. J. Mycobacterium tuberculosis ClpP1 and ClpP2 function together in protein degradation and are required for viability in vitro and during infection. PLoS Pathog. 2012, 8 (2), e1002511.

(39) Ollinger, J.; O'Malley, T.; Kesicki, E. A.; Odingo, J.; Parish, T. Validation of the essential ClpP protease in Mycobacterium tuberculosis as a novel drug target. J. Bacteriol. 2012, 194 (3), 663–668.

(40) Muntel, J.; Fromion, V.; Goelzer, A.; Maa, S.; Mäder, U.; Büttner, K.; Hecker, M.; Becher, D. Comprehensive absolute quantification of the cytosolic proteome of bacillus subtilis by data independent, parallel fragmentation in liquid chromatography/mass spectrometry (LC/MSE). Mol. Cell. Proteomics 2014, 13 (4), 1008–1019.

(41) Gribun, A.; Kimber, M. S.; Ching, R.; Sprangers, R.; Fiebig, K. M.; Houry, W. A. The ClpP double ring tetradecameric protease exhibits plastic ring-ring interactions, and the N termini of its subunits form flexible loops that are essential for ClpXP and ClpAP complex formation. J. Biol. Chem. 2005, 280 (16), 16185–16196.

(42) Yoo, S. J.; Seol, J. H.; Shin, D. H.; Rohrwild, M.; Kang, M. S.; Tanaka, K.; Goldberg, A. L.; Chung, C. H. Purification and characterization of the heat shock proteins HsIV and HsIU that form a new ATP-dependent protease in Escherichia coli. J. Biol. Chem. 1996, 271 (24), 14035–14040.

(43) Gur, E.; Sauer, R. T. Recognition of misfolded proteins by Lon, a AAA+ protease. Genes Dev. 2008, 22 (16), 2267–2277.

(44) Gur, E.; Sauer, R. T. Degrons in protein substrates program the speed and operating efficiency of the AAA+ Lon proteolytic machine. Proc. Natl. Acad. Sci. U. S. A. 2009, 106 (44), 18503–18508.

(45) Krüger, E.; Witt, E.; Ohlmeier, S.; Hanschke, R.; Hecker, M. The Clp proteases of Bacillus subtilis are directly involved in degradation of misfolded proteins. J. Bacteriol. 2000, 182 (11), 3259–3265.

(46) Hughes, A. C.; Subramanian, S.; Dann, C. E.; Kearns, D. B. The C-terminal region of Bacillus subtilis SwrA is required for activity and adaptor-dependent LonA proteolysis. J. Bacteriol. 2018, 200 (6), e00659–17.

(47) Mukherjee, S.; Bree, A. C.; Liu, J.; Patrick, J. E.; Chien, P.; Kearns, D. B. Adaptor-mediated Lon proteolysis restricts Bacillus subtilis hyperflagellation. Proc. Natl. Acad. Sci. U. S. A. 2015, 112 (1), 250–255.

(48) Mielich-Süss, B.; Schneider, J.; Lopez, D. Overproduction of flotillin influences cell differentiation and shape in Bacillus subtilis. MBio 2013, 4 (6), e00719–13.

(49) Thi Nguyen, H. B.; Schumann, W. The sporulation control gene spo0M of Bacillus subtilis is a target of the FtsH metalloprotease. Res. Microbiol. 2012, 163 (2), 114–118.

(50) Bradshaw, N.; Losick, R. Asymmetric division triggers cell-specific gene expression through coupled capture and stabilization of a phosphatase. Elife 2015, 4, e08145.

(51) Koo, B. M.; Kritikos, G.; Farelli, J. D.; Todor, H.; Tong, K.; Kimsey, H.; Wapinski, I.; Galardini, M.; Cabal, A.; Peters, J. M.; et al. Construction and Analysis of Two Genome-Scale Deletion Libraries for Bacillus subtilis. Cell Syst. 2017, 4 (3), 291–305.

(52) Lazar, C.; Gatto, L.; Ferro, M.; Bruley, C.; Burger, T. Accounting for the Multiple Natures of Missing Values in Label-Free Quantitative Proteomics Data Sets to Compare Imputation Strategies. J. Proteome Res. 2016, 15 (4), 1116–1125.

(53) Mi, H.; Muruganujan, A.; Ebert, D.; Huang, X.; Thomas, P. D. PANTHER version 14: More genomes, a new PANTHER GO-slim and improvements in enrichment analysis tools. Nucleic Acids Res. 2019, 47, D419–D426.

(54) Belenky, P.; Ye, J. D.; Porter, C. B. M.; Cohen, N. R.; Lobritz, M. A.; Ferrante, T.; Jain, S.; Korry, B. J.; Schwarz, E. G.; Walker, G. C.; et al. Bactericidal Antibiotics Induce Toxic Metabolic Perturbations that Lead to Cellular Damage. Cell Rep. 2015, 13 (5), 968–980.

(55) Adams, D. W.; Errington, J. Bacterial cell division: Assembly, maintenance and disassembly of the Z ring. Nature Reviews Microbiology. 2009, pp 642–653.

(56) Errington, J.; Wu, L. J. Cell cycle machinery in Bacillus subtilis. Subcell. Biochem. 2017, 84, 67–101.

(57) Eymann, C.; Dreisbach, A.; Albrecht, D.; Bernhardt, J.; Becher, D.; Gentner, S.; Tam, L. T.; Büttner, K.; Buurman, G.; Scharf, C.; et al. A comprehensive proteome map of growing Bacillus subtilis cells. Proteomics 2004, 4, 2849–2876.

(58) Ogino, H.; Wachi, M.; Ishii, A.; Iwai, N.; Nishida, T.; Yamada, S.; Nagai, K.; Sugai, M. FtsZ-dependent localization of GroEL protein at possible division sites. Genes to Cells 2004, 9 (9), 765–771.

(59) Bottomley, A. L.; Liew, A. T. F.; Kusuma, K. D.; Peterson, E.; Seidel, L.; Foster, S. J.; Harry, E. J. Coordination of chromosome segregation and cell division in Staphylococcus aureus. Front. Microbiol. 2017, 8, 1575.

(60) Sugimoto, S.; Saruwatari, K.; Higashi, C.; Sonomoto, K. The proper ratio of GrpE to DnaK is important for protein quality control by the DnaK-DnaJ-GrpE chaperone system and for cell division. Microbiology 2008, 154 (7), 1876–1885.

(61) Sinai, L.; Rosenberg, A.; Segev, E.; Correspondence, B.-Y.; Smith, Y.; Ben-Yehuda, S. The Molecular Timeline of a Reviving Bacterial Spore. Mol. Cell 2015, 57, 695–707.

(62) Vega-Cabrera, L. A.; Guerrero, A.; Rodríguez-Mejía, J. L.; Tabche, M. L.; Wood, C. D.; Gutiérrez-Rios, R. M.; Merino, E.; Pardo-López, L. Analysis of Spo0M function in Bacillus subtilis. PLoS One 2017, 12 (2), e0172737.

(63) Gamba, P.; Veening, J. W.; Saunders, N. J.; Hamoen, L. W.; Daniel, R. A. Two-step assembly dynamics of the Bacillus subtilis divisome. J. Bacteriol. 2009, 191 (13), 4186–4194.

(64) Hammond, L. R.; White, M. L.; Eswara, P. J. ¡vIVA la DivIVA! Journal of bacteriology. NLM (Medline) November 1, 2019, pp e00245–19.

(65) Dajkovic, A.; Lan, G.; Sun, S. X.; Wirtz, D.; Lutkenhaus, J. MinC Spatially Controls Bacterial Cytokinesis by Antagonizing the Scaffolding Function of FtsZ. Curr. Biol. 2008, 18 (4), 235–244.

(66) Scheffers, D. J. The effect of MinC on FtsZ polymerization is pH dependent and can be counteracted by ZapA. FEBS Lett. 2008, 582 (17), 2601–2608.

(67) Blasios, V.; Bisson-Filho, A. W.; Castellen, P.; Nogueira, M. L. C.; Bettini, J.; Portugal, R. V.; Zeri, A. C. M.; Gueiros-Filho, F. J. Genetic and Biochemical Characterization of the MinC-FtsZ Interaction in Bacillus subtilis. PLoS One 2013, 8 (4), e60690.

(68) Marston, A. L.; Errington, J. Selection of the midcell division site in Bacillus subtilis through MinD-dependent polar localization and activation of MinC. Mol. Microbiol. 1999, 33 (1), 84–96.

(69) Eswaramoorthy, P.; Erb, M. L.; Gregory, J. A.; Silverman, J.; Pogliano, K.; Pogliano, J.; Ramamurthi, K. S. Cellular architecture mediates DivIVA ultrastructure and regulates min activity in Bacillus subtilis. MBio 2011, 2 (6), e00257–11.

(70) Briley, K.; Prepiak, P.; Dias, M. J.; Hahn, J.; Dubnau, D. Maf acts downstream of ComGA to arrest cell division in competent cells of B. subtilis. Mol. Microbiol. 2011, 81 (1), 23–39.

(71) Kloosterman, T. G.; Lenarcic, R.; Willis, C. R.; Roberts, D. M.; Hamoen, L. W.; Errington, J.; Wu, L. J. Complex polar machinery required for proper chromosome segregation in vegetative and sporulating cells of Bacillus subtilis. Mol. Microbiol. 2016, 101 (2), 333–350.

(72) Trip, E. N.; Veening, J.; Stewart, E. J.; Errington, J.; Scheffers, D. Balanced transcription of cell division genes in Bacillus subtilis as revealed by single cell analysis. Environ. Microbiol. 2013, 15 (12), 3196–3209.

(73) Jana, B.; Tao, L.; Biswas, I. Strain-Dependent Recognition of a Unique Degradation Motif by ClpXP in Streptococcus mutans. mSphere 2016, 1 (6), e00287–16.

(74) Yu, Y.; Yan, F.; He, Y.; Qin, Y.; Chen, Y.; Chai, Y.; Guo, J. H. The ClpY-ClpQ protease regulates multicellular development in Bacillus subtilis. Microbiology 2018, 164 (5), 848–862.

(75) Levin, P. A.; Margolis, P. S.; Setlow, P.; Losick, R.; Sun, D. Identification of Bacillus subtilis Genes for Septum Placement and Shape Determination. J. Bacteriol. 1992, 174 (21), 6717–6728.

(76) Begg, K.; Nikolaichik, Y.; Crossland, N.; Donachie, W. D. Roles of FtsA and FtsZ in activation of division sites. J. Bacteriol. 1998, 180 (4), 881–884.

(77) Dai, K.; Lutkenhaus, J. The proper ratio of FtsZ to FtsA is required for cell division to occur in Escherichia coli. J. Bacteriol. 1992, 174 (19), 6145–6151.

(78) Dewar, S. J.; Begg, K. J.; Donachie, W. D. Inhibition of cell division initiation by an imbalance in the ratio of FtsA to FtsZ. J. Bacteriol. 1992, 174 (19), 6314–6316.

(79) Ma, X.; Ehrhardt, D. W.; Margolin, W. Colocalization of cell division proteins FtsZ and FtsA to cytoskeletal structures in living Escherichia coli cells by using green fluorescent protein. Proc. Natl. Acad. Sci. U. S. A. 1996, 93 (23), 12998–13003.

(80) Cha, J. H.; Stewart, G. C. The DivIVA minicell locus of Bacillus subtilis. J. Bacteriol. 1997, 179 (5), 1671–1683.

(81) Levin, P. A.; Schwartz, R. L.; Grossman, A. D. Polymer stability plays an important role in the positional regulation of FtsZ. J. Bacteriol. 2001, 183 (18), 5449–5452.

(82) Szwedziak, P.; Löwe, J. Do the divisome and elongasome share a common evolutionary past? Current Opinion in Microbiology. Curr Opin Microbiol December 2013, pp 745–751.

(83) Abhayawardhane, Y.; Stewart, G. C. Bacillus subtilis possesses a second determinant with extensive sequence similarity to the escherichia coli mreB morphogene. J. Bacteriol. 1995, 177 (3), 765–773.

(84) Carballido-López, R.; Formstone, A.; Li, Y.; Ehrlich, S. D.; Noirot, P.; Errington, J. Actin Homolog MreBH Governs Cell Morphogenesis by Localization of the Cell Wall Hydrolase LytE. Dev. Cell 2006, 11 (3), 399–409.

(85) Salje, J.; van den Ent, F.; de Boer, P.; Löwe, J. Direct Membrane Binding by Bacterial Actin MreB. Mol. Cell 2011, 43 (3), 478–487.

(86) Garti-Levi, S.; Hazan, R.; Kain, J.; Fujita, M.; Ben-Yehuda, S. The FtsEX ABC transporter directs cellular differentiation in Bacillus subtilis. Mol. Microbiol. 2008, 69 (4), 1018–1028.

(87) De Leeuw, E.; Graham, B.; Phillips, G. J.; Ten Hagen-Jongman, C. M.; Oudega, B.; Luirink, J. Molecular characterization of Escherichia coli FtsE and FtsX. Mol. Microbiol. 1999, 31 (3), 983–993.

(88) Yamaguchi, H.; Furuhata, K.; Fukushima, T.; Yamamoto, H.; Sekiguchi, J. Characterization of a new Bacillus subtilis peptidoglycan hydrolase gene, yvcE (named cwlO), and the enzymatic properties of its encoded protein. J. Biosci. Bioeng. 2004, 98 (3), 174–181.

(89) Kawai, Y.; Daniel, R. A.; Errington, J. Regulation of cell wall morphogenesis in Bacillus subtilis by recruitment of PBP1 to the MreB helix. Mol. Microbiol. 2009, 71 (5), 1131–1144.

(90) Kawai, Y.; Asai, K.; Errington, J. Partial functional redundancy of MreB isoforms, MreB, Mbl and MreBHp in cell morphogenesis of Bacillus subtilis. Mol. Microbiol. 2009, 73 (4), 719–731.

(91) Wachi, M.; Matsuhashi, M. Negative control of cell division by mreB, a gene that functions in determining the rod shape of Escherichia coli cells. J. Bacteriol. 1989, 171 (6), 3123–3127.

(92) Ranjit, D. K.; Liechti, G. W.; Maurelli, A. T. Chlamydial MreB directs cell division and peptidoglycan synthesis in Escherichia coli in the absence of FtsZ activity. MBio 2020, 11 (1).

(93) Schirner, K.; Errington, J. Influence of heterologous MreB proteins on cell morphology of Bacillus subtilis. Microbiology 2009, 155 (11), 3611–3621.

(94) Brunet, Y. R.; Wang, X.; Rudner, D. Z. SweC and SweD are essential co-factors of the FtsEX-CwlO cell wall hydrolase complex in Bacillus subtilis. PLoS Genet. 2019, 15 (8), e1008296.

(95) Do, T.; Page, J. E.; Walker, S. Uncovering the activities, biological roles, and regulation of bacterial cell wall hydrolases and tailoring enzymes. Journal of Biological Chemistry. American Society for Biochemistry and Molecular Biology Inc. March 6, 2020, pp 3347–3361.

(96) Molle, V.; Fujita, M.; Jensen, S. T.; Eichenberger, P.; González-Pastor, J. E.; Liu, J. S.; Losick, R. The Spo0A regulon of Bacillus subtilis. Mol. Microbiol. 2003, 50 (5), 1683–1701.

(97) Al-Karadaghi, S.; Hansson, M.; Nikonov, S.; Jönsson, B.; Hederstedt, L. Crystal structure of ferrochelatase: the terminal enzyme in heme biosynthesis. Structure 1997, 5 (11), 1501–1510.

(98) Klaeger, S.; Gohlke, B.; Perrin, J.; Gupta, V.; Heinzlmeir, S.; Helm, D.; Qiao, H.; Bergamini, G.; Handa, H.; Savitski, M. M.; et al. Chemical proteomics reveals ferrochelatase as a common off-target of kinase inhibitors. ACS Chem. Biol. 2016, 11 (5), 1245–1254.

(99) Kahl, B. C.; Becker, K.; Löffler, B. Clinical significance and pathogenesis of staphylococcal small colony variants in persistent infections. Clinical Microbiology Reviews. American Society for Microbiology March 9, 2016, pp 401–427.

(100) Potter, K. D.; Merlino, N. M.; Jacobs, T.; Gollnick, P. TRAP binding to the Bacillus subtilis trp leader region RNA causes efficient transcription termination at a weak intrinsic terminator. Nucleic Acids Res. 2011, 39 (6), 2092–2102.

(101) Yang, W.-J.; Yanofsky, C. Effects of Tryptophan Starvation on Levels of the trp RNA-Binding Attenuation Protein (TRAP) and Anti-TRAP Regulatory Protein and Their Influence on trp Operon Expression in Bacillus subtilis. J. Bacteriol. 2005, 187 (6), 1884–1891.

(102) Hobley, L.; Ostrowski, A.; Rao, F. V.; Bromley, K. M.; Porter, M.; Prescott, A. R.; MacPhee, C. E.; Van Aalten, D. M. F.; Stanley-Wall, N. R. BslA is a self-assembling bacterial hydrophobin that coats the Bacillus subtilis biofilm. Proc. Natl. Acad. Sci. U. S. A. 2013, 110 (33), 13600–13605.

(103) Lesniak, J.; Barton, W. A.; Nikolov, D. B. Structural and functional features of the *Escherichia coli* hydroperoxide resistance protein OsmC. Protein Sci. 2003, 12 (12), 2838–2843.

(104) Auger, S.; Gomez, M. P.; Danchin, A.; Martin-Verstraete, I. The PatB protein of Bacillus subtilis is a C-S-lyase. Biochimie 2005, 87 (2), 231–238.

(105) Gentry-Weeks, C. R.; Spokes, J.; Thompson, J. β-Cystathionase from *Bordetella avium*. J. Biol. Chem. 1995, 270 (13), 7695–7702.

(106) Ejim, L. J.; Blanchard, J. E.; Koteva, K. P.; Sumerfield, R.; Elowe, N. H.; Chechetto, J. D.; Brown, E. D.; Junop, M. S.; Wright, G. D. Inhibitors of bacterial cystathionine β-lyase: leads for new antimicrobial agents and probes of enzyme structure and function. J. Med. Chem. 2007, 50 (4), 755–764.

(107) Bradshaw, J. M.; McFarland, J. M.; Paavilainen, V. O.; Bisconte, A.; Tam, D.; Phan, V. T.; Romanov, S.; Finkle, D.; Shu, J.; Patel, V.; et al. Prolonged and tunable residence time using reversible covalent kinase inhibitors. Nat. Chem. Biol. 2015, 11 (7), 525–531.

(108) Dornan, M. H.; Krishnan, R.; MacKlin, A. M.; Selman, M.; El Sayes, N.; Son, H. H.; Davis, C.; Chen, A.; Keillor, K.; Le, P. J.; et al. First-in-class small molecule potentiators of cancer virotherapy. Sci. Rep. 2016, 6, 26786.

(109) Shajani, Z.; Sykes, M. T.; Williamson, J. R. Assembly of bacterial ribosomes. Annu. Rev. Biochem. 2011, 80 (1), 501–526.

(110) Corrigan, R. M.; Bellows, L. E.; Wood, A.; Gründling, A. PpGpp negatively impacts ribosome assembly affecting growth and antimicrobial tolerance in Gram-positive bacteria. Proc. Natl. Acad. Sci. 2016, 113 (12), E1710–E1719.

(111) Irving, S. E.; Corrigan, R. M. Triggering the stringent response: signals responsible for activating (p)ppGpp synthesis in bacteria. Microbiology 2018, 164 (3), 268–276.

(112) Starosta, A. L.; Lassak, J.; Jung, K.; Wilson, D. N. The bacterial translation stress response. FEMS Microbiol. Rev. 2014, 38 (6), 1172–1201.

(113) Ollinger, J.; Song, K. B.; Antelmann, H.; Hecker, M.; Helmann, J. D. Role of the Fur regulon in iron transport in Bacillus subtilis. J. Bacteriol. 2006, 188 (10), 3664–3673.

(114) Miethke, M.; Westers, H.; Blom, E.-J.; Kuipers, O. P.; Marahiel, M. A. Iron starvation triggers the stringent response and induces amino acid biosynthesis for bacillibactin production in Bacillus subtilis. J. Bacteriol. 2006, 188 (24), 8655–8657.

(115) Haranahalli, K.; Tong, S.; Ojima, I. Recent advances in the discovery and development of antibacterial agents targeting the cell-division protein FtsZ. Bioorg. Med. Chem 2016, 24 (24), 6354–6369.

(116) Stokes, N. R.; Baker, N.; Bennett, J. M.; Collins, I.; Czaplewski, L. G.; Logan, A.; Macdonald, R.; Macleod, L.; Peasley, H.; Mitchell, J. P.; et al. An Improved Small-Molecule Inhibitor of FtsZ with Superior In Vitro Potency, Drug-Like Properties, and In Vivo Efficacy. Antimicrob. Agents Chemother. 2013, 57 (1), 317–325.

(117) Typas, A.; Banzhaf, M.; Gross, C. A.; Vollmer, W. From the regulation of peptidoglycan synthesis to bacterial growth and morphology. Nature Reviews Microbiology. Nature Publishing Group February 28, 2012, pp 123–136.

(118) Zhao, H.; Patel, V.; Helmann, J. D.; Dörr, T. Don't let sleeping dogmas lie: new views of peptidoglycan synthesis and its regulation. Mol. Microbiol. 2017, 106 (6), 847–860.

(119) Meng, L.; Mohan, R.; Kwok, B. H. B.; Elofsson, M.; Sin, N.; Crews, C. M. Epoxomicin, a potent and selective proteasome inhibitor, exhibits in vivo antiinflammatory activity. Proc. Natl. Acad. Sci. 1999, 96 (18), 10403–10408.

(120) Groll, M.; Kim, K. B.; Kairies, N.; Huber, R.; Crews, C. M. Crystal Structure of Epoxomicin: 20S Proteasome Reveals a Molecular Basis for Selectivity of α',β'-Epoxyketone Proteasome Inhibitors. J. Am. Chem. Soc. 2000, 122 (6), 1237–1238.

(121) Feling, R. H.; Buchanan, G. O.; Mincer, T. J.; Kauffman, C. A.; Jensen, P. R.; Fenical, W. Salinosporamide A: A Highly Cytotoxic Proteasome Inhibitor from a Novel Microbial Source, a Marine Bacterium of the New Genus Salinospora. Angew. Chemie Int. Ed. 2003, 42 (3), 355–357.

(122) Borissenko, L.; Groll, M. 20S proteasome and its inhibitors: crystallographic knowledge for drug development. Chem. Rev. 2007, 107 (3), 687–717.

(123) Malik, I. T.; Brötz-Oesterhelt, H. Conformational control of the bacterial Clp protease by natural product antibiotics. Nat. Prod. Rep. 2017, 34 (7), 815–831.

(124) Kirstein, J.; Hoffmann, A.; Lilie, H.; Schmidt, R.; Helga, R. W.; Heike, B. O.; Mogk, A.; Turgay, K. The antibiotic ADEP reprogrammes ClpP, switching it from a regulated to an uncontrolled protease. EMBO Mol. Med. 2009, 1 (1), 37–49.

(125) Vasudevan, D.; Rao, S. P. S.; Noble, C. G. Structural basis of mycobacterial inhibition by Cyclomarin A. J. Biol. Chem. 2013, 288 (43), 30883–30891.

(126) Gavrish, E.; Sit, C. S.; Cao, S.; Kandror, O.; Spoering, A.; Peoples, A.; Ling, L.; Fetterman, A.; Hughes, D.; Bissell, A.; et al. Lassomycin, a ribosomally synthesized cyclic peptide, kills Mycobacterium tuberculosis by targeting the ATP-dependent protease ClpC1P1P. Chem. Biol. 2014, 21 (4), 509–518.

(127) Gao, W.; Kim, J.-Y.; Anderson, J. R.; Akopian, T.; Hong, S.; Jin, Y.-Y.; Kandror, O.; Kim, J.-W.; Lee, I.-A.; Lee, S.-Y.; et al. The cyclic peptide ecumicin targeting ClpC1 is active against Mycobacterium tuberculosis in vivo. Antimicrob. Agents Chemother. 2015, 59 (2), 880–889.

(128) Maurizis, M. R.; Clark, W. P.; Kim, S. H.; Gottesman, S. Clp P represents a unique family of serine proteases. J. Biol. Chem. 1990, 265 (21), 12546–12552.

(129) Wang, J.; Hartling, J. A.; Flanagan, J. M. The structure of ClpP at 2.3 Å resolution suggests a model for ATP-dependent proteolysis. Cell 1997, 91 (4), 447–456.

(130) Valas, R. E.; Bourne, P. E. Rethinking proteasome evolution: Two novel bacterial proteasomes. J. Mol. Evol. 2008, 66 (5), 494–504.

(131) Maurizi, M. R.; Xia, D. Protein binding and disruption by Clp/Hsp100 chaperones. Structure 2004, 12 (2), 175–183.

(132) Schirmer, E. C.; Glover, J. R.; Singer, M. A.; Lindquist, S. HSP100/Clp proteins: a common mechanism explains diverse functions. Trends Biochem. Sci. 1996, 21 (8), 289–296.

(133) Culp, E.; Wright, G. D. Bacterial proteases, untapped antimicrobial drug targets. Journal of Antibiotics. Nature Publishing Group April 1, 2017, pp 366–377.

(134) Moreno-Cinos, C.; Goossens, K.; Salado, I. G.; Van Der Veken, P.; De Winter, H.; Augustyns, K. ClpP protease, a promising antimicrobial target. International Journal of Molecular Sciences. MDPI AG May 1, 2019, p 2232.

(135) Böttcher, T.; Sieber, S. A. β-Lactones as privileged structures for the active-site labeling of versatile bacterial enzyme classes. Angew. Chemie Int. Ed. 2008, 47 (24), 4600–4603.

(136) Hackl, M. W.; Lakemeyer, M.; Dahmen, M.; Glaser, M.; Pahl, A.; Lorenz-Baath, K.; Menzel, T.; Sievers, S.; Böttcher, T.; Antes, I.; et al. Phenyl esters are potent inhibitors of caseinolytic protease P and reveal a stereogenic switch for deoligomerization. J. Am. Chem. Soc. 2015, 137 (26), 8475–8483.

(137) Ju, Y.; He, L.; Zhou, Y.; Yang, T.; Sun, K.; Song, R.; Yang, Y.; Li, C.; Sang, Z.; Bao, R.; et al. Discovery of novel peptidomimetic boronate ClpP inhibitors with noncanonical enzyme mechanism as potent virulence blockers in vitro and in vivo. J. Med. Chem. 2020, 63 (6), 3104–3119.

(138) Compton, C. L.; Schmitz, K. R.; Sauer, R. T.; Sello, J. K. Antibacterial activity of and resistance to small molecule inhibitors of the clpp peptidase. ACS Chem. Biol. 2013, 8 (12), 2669–2677.

(139) Shoji, M.; Cui, L.; Iizuka, R.; Komoto, A.; Neoh, H. M.; Watanabe, Y.; Hishinuma, T.; Hiramatsu, K. WalK and clpP mutations confer reduced vancomycin susceptibility in Staphylococcus aureus. Antimicrob. Agents Chemother. 2011, 55 (8), 3870–3881.

(140) Song, Y.; Rubio, A.; Jayaswal, R. K.; Silverman, J. A.; Wilkinson, B. J. Additional routes to Staphylococcus aureus daptomycin resistance as revealed by comparative genome sequencing, transcriptional profiling, and phenotypic studies. PLoS One 2013, 8 (3).

(141) Gominet, M.; Seghezzi, N.; Mazodier, P. Acyl depsipeptide (ADEP) resistance in streptomyces. Microbiology 2011, 157 (8), 2226–2234.

(142) Brötz-Oesterhelt, H.; Beyer, D.; Kroll, H. P.; Endermann, R.; Ladel, C.; Schroeder, W.; Hinzen, B.; Raddatz, S.; Paulsen, H.; Henninger, K.; et al. Dysregulation of bacterial proteolytic machinery by a new class of antibiotics. Nat. Med. 2005, 11 (10), 1082–1087.

(143) Oldfield, E.; Feng, X. Resistance-resistant antibiotics. Trends in Pharmacological Sciences. Elsevier Ltd 2014, pp 664–674.

(144) Gray, D. A.; Wenzel, M. Multitarget approaches against multiresistant superbugs. ACS Infect. Dis. 2020, 6 (6), 1346–1365.

