## Supplemental figures and methods for "Armeniaspirols inhibit the AAA+ proteases ClpXP and ClpYQ leading to cell division arrest in Gram-positive bacteria"

### Experimental Procedures

#### Cloning and protein purification

*B. subtilis clp* genes were PCR amplified from *B. subtilis* 168 genomic DNA (Promega Wizard Genomic DNA Purification Kit). The clpQ gene was cloned into a pET21 vector (C-terminal His tag, AmpR). The resulting plasmid (pPL29) was transformed into chemically competent *E. coli* BL21(DE3) for protein expression. 400 mL LB media was inoculated with 0.5% (*v/v*) of an overnight pre-culture. The culture was grown at 37°C (200 rpm) and expression was induced with 1 mM isopropyl-1-thio-β-D-galactopyranoside at an optical density ($OD_{600}$) of 0.5. The culture was grown at 37°C (200 rpm) for 4 h. The cells were pelleted at 4,000 rpm for 20 min and re-suspended in lysis buffer (50 mM Tris, 100 mM NaCl, 1 µg/mL leupeptin, 1 µg/mL pepstatin A, 1 mg/mL lysozyme, 10% glycerol, pH 8.0). Mechanical cell lysis was achieved by 30 s sonication. The cell debris was pelleted at 10,000 rpm for 60 min and the supernatant was incubated with 800 µL 50% Ni-NTA agarose resin (QIAGEN) for 30 min at 4°C with gentle shaking. The lysate was loaded onto a column and the flow-through was collected. The resin was washed sequentially with 5 mL elution buffer (100 mM Tris, 300 mM NaCl, pH 8.0) containing 0 mM, 20 mM, 100 mM and 250 mM imidazole. Fractions were analyzed by SDS-PAGE (4-20% Mini-PROTEAN TGX Precast Gels; Bio-Rad). Fractions containing purified protein were pooled and buffer exchanged (50 mM Tris, 250 mM NaCl, pH 8.0) using a 10 kDa Amicon Ultra-15 centrifugal filter unit (Millipore Sigma). The concentrated protein was stored at -80°C with 30% (w/w) glycerol. The *B. subtilis clpY* gene was cloned into pET28b (N-terminal His tag, KanR), and the resulting plasmid (pPL31) was purified as above. All protein concentrations were determined by Bradford assay.

The *E. coli clpX* plasmid (pProEx-Htb-ClpX) was expressed as above in 0.8 L LB media. The cells were pelleted at 4,000 rpm for 20 min and re-suspended in 20 mL lysis buffer (25 mM Tris, 500 mM NaCl, 10 mM imidazole, 1 µg/mL leupeptin, 1 µg/mL pepstatin A, 1 mg/mL lysozyme, 10% (w/w) glycerol, pH 8.0). Mechanical cell lysis was achieved by 30 s sonication. The cell debris was pelleted at 10,000 rpm for 60 min and the supernatant was incubated with 2.4 mL 50% Ni-NTA agarose resin (QIAGEN) for 30 min at 4°C with gentle shaking. The lysate was loaded onto a column and the flow-through was collected. The resin was washed 2X with 6 mL lysis buffer (25 mM Tris, 500 mM NaCl, 10 mM imidazole, 10% (w/w) glycerol, pH 8.0), 3X with 10 mL wash buffer (25 mM Tris, 500 mM NaCl, 50 mM imidazole, pH 8.0), and 2X with 5 mL elution buffer (25 mM Tris, 500 mM NaCl, 250 mM imidazole, pH 8.0). Fractions were analyzed by SDS-PAGE (4-20% Mini-PROTEAN TGX Precast Gels; Bio-Rad). Fractions containing purified protein were pooled and buffer exchanged (50 mM Tris, 200 mM KCl, 25 mM $MgCl_2$, 1 mM DTT, 0.1 mM EDTA, 10% (w/w) glycerol pH 8.0) overnight at 4°C. The concentrated protein was stored in small aliquots at -80°C.

The *E. coli clpP* plasmid (pET9a-ClpP) was expressed as above in 0.8 L LB media. The cells were pelleted at 4,000 rpm for 20 min and re-suspended in lysis buffer (50 mM Tris, 150 mM KCl, 1 µg/mL leupeptin, 1 µg/mL pepstatin A, 1 mg/mL lysozyme, 10% (w/w) glycerol, pH 8.0). Mechanical cell lysis was achieved by 60 s sonication. The cell debris was pelleted at 10,000 rpm for 60 min and saturated ammonium sulfate was added to the supernatant slowly with stirring until 40% saturation. Stirring was continued for 60 min at 4°C. The lysate was centrifuged at 10,000 rpm for 30 min and the pellet was discarded. Saturated ammonium sulfate was added further until 60% saturation. Stirring was continued for 60 min at 4°C. The lysate was centrifuged at 10,000 rpm for 30 min and the pellet was discarded. The lysate was buffer exchanged into Buffer A (50 mM Tris, 150 mM KCl, 1 mM DTT, 10% (w/w) glycerol, pH 8.0) and filtered through a 0.2 µm syringe filter. The solution was injected onto a 1 mL Q sepharose column (GE Healthcare) and eluted with a step gradient of 0%, 10%, 15%, 20%, 25%, 30% and 50% Buffer B (50 mM Tris, 1 M KCl, 1 mM DTT, 10% (w/w) glycerol, pH 8.0) each over 1.5 column volumes. 0.5 mL fractions were collected and analyzed by SDS-PAGE. ClpP containing fractions were pooled and buffer exchanged into 50 mM MES pH 6.0 using a 10 kDa Amicon Ultra-15 centrifugal filter unit (Millipore Sigma). The solution was injected onto a 1 mL SP sepharose column (GE Healthcare) and eluted with a step gradient of 0%, 5%, 10%, 15%, 20%, 25% and 30% Buffer C (50 mM MES, 1 M KCl, 10% (w/w) glycerol, pH 6.0) each over 2 column volumes. 0.5 mL fractions were collected and analyzed by SDS-PAGE. Fractions containing ClpP were concentrated using a 10 kDa Amicon Ultra-15 centrifugal filter unit (Millipore Sigma). The concentrated protein was stored at -80°C with 30% (w/w) glycerol.

#### Peptide hydrolysis assay

Peptide hydrolysis was assayed using the Cbz-Gly-Gly-Leu-AMC (Millipore Sigma) substrate. 0.1 mL reaction assays were done in Nunc 96-well microplates for fluorescence-based assays (ThermoFisher Scientific). Assays were composed of purified bsClpQ and bsClpY protein, 0.1 M Tris (pH 8.0), 0.1 mM Cbz-GGL-AMC, 10 mM $MgCl_2$, 1 mM ATP, 1 mM TCEP, and 1 mM EDTA. Peptide hydrolysis for ecClpX and ecClpP was monitored with the substrate Suc-Leu-Tyr-AMC (Millipore Sigma) under the same conditions. A continuous assay of AMC release was monitored at 37°C using a Synergy H4 microplate reader (BioTek). Excitation and emission for Cbz-GGL-AMC were measured at 355 nm and 460 nm, respectively. Excitation and emission for Suc-LY-AMC were measured at 360 nm and 440 nm, respectively. Inhibition was observed with varying concentrations of **1**.

**ATP hydrolysis assay**

ATP hydrolysis was monitored by a discontinuous assay with a malachite green colour reagent. 3 volumes 0.045% (w/v) malachite green in $dH_2O$ and 1 volume 4.2% (w/v) ammonium molybdate in 4 M HCl were mixed to make the colour reagent. After 20 min of shaking at room temperature in the dark, 100 µL of 2% (w/v) Triton X-100 per 5 mL colour reagent was added, and the resulting solution was filtered through a 0.2 µm syringe filter. Reaction assays contained purified bsClpY and bsClpQ, 1 mM ATP, 10 mM $MgCl_2$, 1 mM TCEP, 1 mM EDTA, 0.1 M Tris (pH 8.0). At specified time intervals, 25 µL of the assay was added to 100 µL of the colour reagent in a 96-well plate. The mixture was incubated at room temperature for 5 min and the absorbance ($A_{650}$) was read using a Synergy H4 microplate reader (BioTek). Inhibition was observed with varying concentrations of **1**.

**Minimum inhibitory concentration**

MICs were carried out in Mueller-Hinton broth, 100 µL assays in 96-well plates. Sequential concentrations of compound were pipetted into each column through two-fold serial dilutions. 5 µL of bacterial culture ($OD_{600}$ 0.07-0.1) was inoculated into each well and incubated at 37°C for 16 h. Growth was observed by $OD_{600}$ readings using a Synergy H4 microplate reader (BioTek). The MIC was determined by the lowest concentration of compound that prevented bacterial growth.

**Microscopy and image analysis**

Biological triplicates of *B. subtilis* 168 was grown in Mueller-Hinton broth with (1 µg/ml) and without compound **1** until an $OD_{600}$ of 0.2-0.25 was reached. 1 mL bacterial culture was prepared on #1.5 coverslips (ThermoFisher Scientific) washed with poly-lysine(K) in phosphate-buffered saline (PBS). The bacterial dilution was fixed with 4% paraformaldehyde-PBS for 10 min at room temperature, and subsequently washed with filter-sterilized 100 mM glycine-PBS for 5 min. The membrane was stained with 5 µg/ml FM 4-64FX (Invitrogen) fluorescent dye while the nucleoid was stained with 2 µg/mL DAPI (Millipore Sigma) fluorescent dye. Coverslips were mounted on Superfrost Plus microscope slides (ThermoFisher Scientific) with Mowiol mounting medium. The slides were left to dry overnight in the dark. Images were obtained with an LSM 880 confocal microscope (Zeiss) using a 100X/1.4 oil immersion objective. Image analysis and cell length measurements were conducted using ZEN Blue (Zeiss) software.

**Whole proteome extraction and dimethyl labeling**

*B. subtilis* 168 was grown in 5 mL Mueller-Hinton broth overnight at 37°C. The culture was diluted 1/50 in fresh 50 mL Mueller-Hinton broth with (1 µg/mL) or without compound **1** and incubated for 6 hours at 37°C (200 rpm). The cells were pelleted at 4,000 rpm for 30 min and re-suspended in 1.25 mL lysis buffer (50 mM Tris, 100 mM NaCl, 1 µg/mL leupeptin, 1 µg/mL pepstatin A, 1 mg/mL lysozyme, pH 8.0). Mechanical cell lysis was achieved by 30 s sonication. The cell debris was pelleted at 10,000 rpm for 10 min. Protein content was quantified by Bradford assay. 50 µg were transferred to a 1.5 mL microcentrifuge tube and the volume was adjusted to 100 µL with 100 mM TEAB buffer. 5 µL of 200 mM TCEP was added to the mixture and incubated at 55°C for 1 h. 5 µL of 375 mM iodoacetamide was added and further incubated at room temperature for 30 min in the dark. A methanol/chloroform precipitation was performed to obtain protein pellet. The pellet was re-suspended in 100 µL of 50 mM TEAB buffer containing 0.015% Triton X-100. Protein was digested with trypsin overnight at 37°C. A pellet was obtained by vacuum centrifugation and re-suspended with 100 µL of 100 mM TEAB buffer. 4 µL of 4% (*v/v*) formaldehyde was added to the control reaction, while deuterated formaldehyde was added to **1**-containing reactions. 4 µL of 0.6 M $NaBH_3CN$ was added to all samples and incubated in a fumehood for 1 h with rotation. 16 µL of 1% (v/v) ammonia solution was added to quench the reaction. 8 µL of 5% formic acid (FA) was added and differentially labeled samples were mixed 1:1. All samples were desalted with Pierce C18 Tips (ThermoFisher Scientific) prior to LC-MS/MS analysis.

***In vitro* labeling and inhibitor covalent capture**

*In vitro labeling and click reaction*. Fresh *B. subtilis* 168 cell lysate (2 mg/mL) was incubated in 1 mL (2 x 0.5 mL) PBS containing 0.05% Triton X-100 with 1 mM of compound **1** for 2 h, followed by incubation with 20 µM of compound **2** for an additional 2 h. 500 µL of freshly prepared click chemistry mix (100 µM biotin-azide, 2 mM TCEP, 200 µM TBTA, and 2 mM $CuSO_4$) in PBS was added to each probe-labeled lysate and incubated for 2 h at room temperature with gentle shaking. Respective 0.5 mL reactions were combined and quenched by addition of 5 mL acetone and stored at -80 $^{o}$C overnight. The solution was centrifuged at maximum speed for 15 min at 4$^{o}$C. The resulting supernatant was discarded, and the pellet was resuspended with 750 µL cold methanol and sonicated for 5 one-second pulses (30% amplitude). The solution was centrifuged for 5 min at 6,500 g at 4°C. The supernatant was discarded, and this process was repeated an additional 2X. 650 µL of 2.5% SDS in PBS was added to the pellet and sonicated for 15 one-second pulses (30% amplitude). Samples were heated for 5 min at 60°C and centrifuged for 4 min at 6,500 g. The supernatant was transferred to a 15 mL falcon tube and the volume was brought to 8 mL with PBS.

*Streptavidin enrichment*. 100 µL of 50% streptavidin-agarose beads were 3X times with 700 µL PBS in a Bio-Spin chromatography column (Bio-Rad) at 1,000g. The beads were transferred to probe-labeled cell lysates using 500 µL PBS and incubated for 90 min at room temperature. Beads were pelleted for 2 min at 1,400 g at room temperature. 500 µL of supernatant was used to transfer beads to Bio-Spin columns. Beads were washed 3X with 1% SDS and 3X with fresh 6 M urea by short-spin centrifugation (1,000 g maximum speed).

*On-bead digestion*. Beads were washed 1X with PBS, 5X with 50 mM ammonium bicarbonate (ABC), and transferred to a 1.5 mL microcentrifuge tube with 700 µL ABC. Beads were pelleted for 2 min at 1,400 g and the supernatant was discarded. 500 µL of 50 mM ABC with 10 mM DTT was added to samples and heated for 15 min at 65$^{o}$C. 25 µL of 500 mM iodoacetamide was added and lysates were incubated at room temperature in the dark for 30 min. Samples were centrifuged for 2 min at 1,400 rpm and the supernatant was discarded. Beads were washed 3X with 500 µL TEAB (pH 8.5) and resuspended with 100 µL TEAB. 2 µL of 0.5 mg/mL trypsin was added to samples and were rotated at 37$^{o}$C overnight. The samples were pelleted, and the supernatant was centrifuged for 1 min at 1,000 g in new Bio-Spin columns. The flow-through was collected, desalted using Pierce C18 Tips (ThermoFisher Scientific), and dried by vacuum centrifugation prior to LC-MS/MS analysis.

**Mass spectrometry**

The LC-MS/MS system consisted of an Ultimate 3000 nanoRLSC system (Dionex) coupled with an Orbitrap Fusion mass spectrometer (ThermoFisher Scientific) operated in positive ion mode. A 70 µm × 150 mm Luna C18(2) reverse phase column (3 µm; 100-Å pore size; Phenomenex) was packed in-house. The mobile phases consisted of 0.1% (v/v) formic acid in water as buffer A and 0.1% (v/v) formic acid in acetonitrile as buffer B. The sample was loaded onto the column using 2% buffer B at a flow rate of 0.30 µL/min for 105 minutes. The method was designed as follows: a gradient from 2% to 38% buffer B was performed for 70 minutes, a gradient from 38% to 98% buffer B for 9 minutes, 98% buffer B for 10 minutes, gradient from 98% to 2% buffer B for 3 minutes, concluded by a 2% buffer B wash for 10 minutes. The Orbitrap Fusion was run in top speed mode and the MS method consisted of a full MS scan from 350 - 2000 *m/z* with R = 60, 000. Precursor ions were filtered according to monoisotopic precursor selection, +2 to +7 charge state, and 30 seconds ± 10 ppm dynamic exclusion. Fragmentation was performed with collision-induced dissociation (CID) in the linear ion trap. Precursors were isolated using a 2 *m/z* isolation window, and fragmented with a normalized collision energy of 35%.

*Dimethyl labeling processing*. Proteome Discoverer 2.1 (ThermoFisher Scientific) was used for protein identification and quantification. The precursor mass tolerance was set at 10 ppm, and 0.6 Da mass tolerance for fragment ions. Search engine SEQUEST-HT implemented in Proteome Discovery was applied for all MS raw files. Search parameters were set to allow for dynamic modification of methionine oxidation, static modification of cysteine carbamidomethylation, and dimethyl modifications on lysine and N-terminus (light and medium labels). The raw files were searched separately with “light” or “medium” labels in the same workflow. The search database consisted of nonredundant *B. subtilis* protein sequences in FASTA file format from the UniProt/SwissProt database. The FDR was set to 0.01, 0.05, and 0.10 for high, medium, and low protein identifications, respectively. Quantification of peptides and proteins was performed using standard settings provided by Proteome Discoverer.

*Inhibitor covalent capture processing*. Proteome Discoverer 2.1 (ThermoFisher Scientific) was used for protein identification. The precursor mass tolerance was set at 10 ppm, and 0.6 Da mass tolerance for fragment ions. Search engine SEQUEST-HT implemented in Proteome Discovery was applied for all MS raw files. Search parameters were set to allow for dynamic modification of methionine oxidation and static modification of cysteine carbamidomethylation. The search database consisted of nonredundant *B. subtilis* protein sequences in FASTA file format from the UniProt/SwissProt database. The FDR was set to 0.05 for both peptide and protein identifications.

*Proteomics data analysis.* Proteins that were sparsely quantified were removed by a comprehensive data filtering. Only proteins with a FDR confidence of high and 2 or more peptides detected were used in the data analysis. Of these only protein hits that were detected in two of three or three of four replicates in either the treatment or control group were used for the analysis. The values were Log(2) transformed and each replicate dataset was normalized by subtracting the median of the Log(2) transformed distribution form all values. Missing values were imputed from a Gaussian distribution downshift 1.8 σ from the mean with a standard deviation of 0.3 σ.

**GO analysis**

The PANTHER Overrepresentation Test was used to search proteomic data against GO Ontology database (Released 2019-02-02) to identify GO annotations overrepresented in our data when compared to the reference *Bacillus subtilis* genome. Fisher's Exact test with a false discovery rate multiple test correction was used to evaluate the significance.

### Supplementary Figures

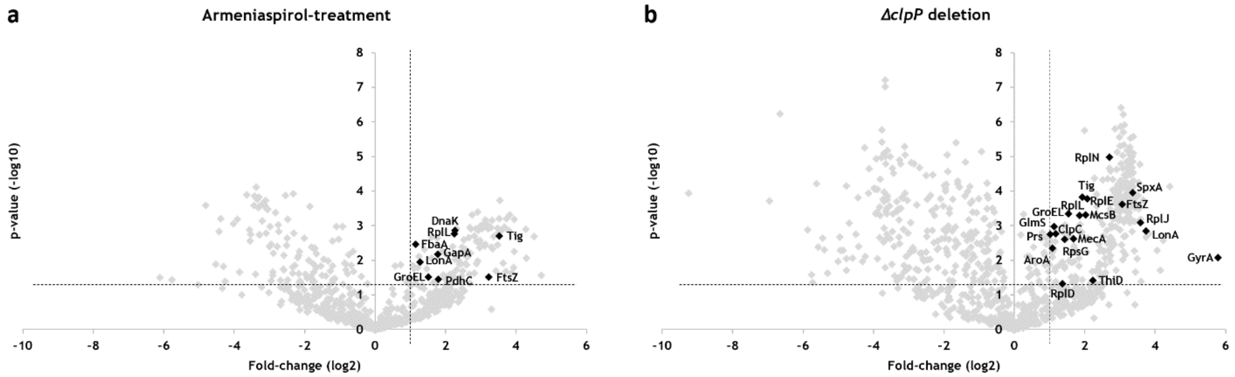


| **ClpP Substrates** | | **Armeniaspirol-treatment** | | ***clpP* deletion** | |
|---|---|---|---|---|---|
| *Accession* | *Description* | *fold-change* | *p-value* | *fold-change* | *p-value* |
| P80698 | Trigger factor, Tig | 11.539 | 0.002 | 3.802 | 0.0001 |
| P17865 | Cell division protein, FtsZ | 9.351 | 0.031 | 8.324 | 0.0002 |
| P02394 | 50S ribosomal protein L7/L12, RplL | 4.732 | 0.002 | 3.601 | 0.0005 |
| P28598 | 60 kDa chaperonin, GroEL | 2.860 | 0.030 | 2.906 | 0.0005 |
| P37945 | Lon protease, LonA | 2.408 | 0.011 | 13.409 | 0.0014 |
| P17820 | Chaperone protein, DnaK | 4.795 | 0.001 | *n.d.* | |
| P21883 | Dihydrolipoyllysine-residue acetyltransferase component of pyruvate dehydrogenase complex, PdhC | 3.468 | 0.035 | *n.d.* | |
| P09124 | Glyceraldehyde-3-phosphate dehydrogenase, GapA | 3.441 | 0.007 | *n.d.* | |
| P13243 | Probable fructose-bisphosphate aldolase, FbaA | 2.212 | 0.003 | *n.d.* | |
| P05653 | DNA gyrase subunit A, GyrA | *n.d.* | | 54.839 | 0.0085 |
| P42923 | 50S ribosomal protein L10, RplJ | *n.d.* | | 12.007 | 0.0008 |
| O31602 | Regulatory protein Spx, SpxA | *n.d.* | | 10.222 | 0.0001 |
| P12875 | 50S ribosomal protein L14, RplN | *n.d.* | | 6.521 | 1.07E-05 |
| P39610 | Pyridoxine kinase. ThiD | *n.d.* | | 4.691 | 0.0380 |
| P12877 | 50S ribosomal protein L5, RplE | *n.d.* | | 4.219 | 0.0001 |
| P37570 | Protein-arginine kinase, McsB | *n.d.* | | 4.068 | 0.0005 |
| P37958 | Adapter protein, MecA | *n.d.* | | 3.224 | 0.0024 |
| P21469 | 30S ribosomal protein S7, RpsG | *n.d.* | | 2.705 | 0.0026 |
| P42921 | 50S ribosomal protein L4, RplD | *n.d.* | | 2.590 | 0.0479 |
| P37571 | Negative regulator of genetic competence, ClpC | *n.d.* | | 2.254 | 0.0017 |
| P0CI73 | Glutamine--fructose-6-phosphate aminotransferase [isomerizing], GlmS | *n.d.* | | 2.181 | 0.0011 |
| P39912 | Protein, AroA | *n.d.* | | 2.116 | 0.0046 |
| P14193 | Ribose-phosphate pyrophosphokinase, Prs | *n.d.* | | 2.019 | 0.0018 |

**Supplementary Figure 1.** Enrichment of several known ClpXP substrates in (a) **1**-treated and (b) *ΔclpP B. subtilis,* supporting a mechanism involving inhibition of ClpXP. Protein fold-change and p-values are identified for proteins known to be ClpP substrates (*n.d.*; not detected). Significance is defined as fold-change > 2 and p-value < 0.05. Data is shown in graphical and tabular format and MS analysis was performed as described in Methods.

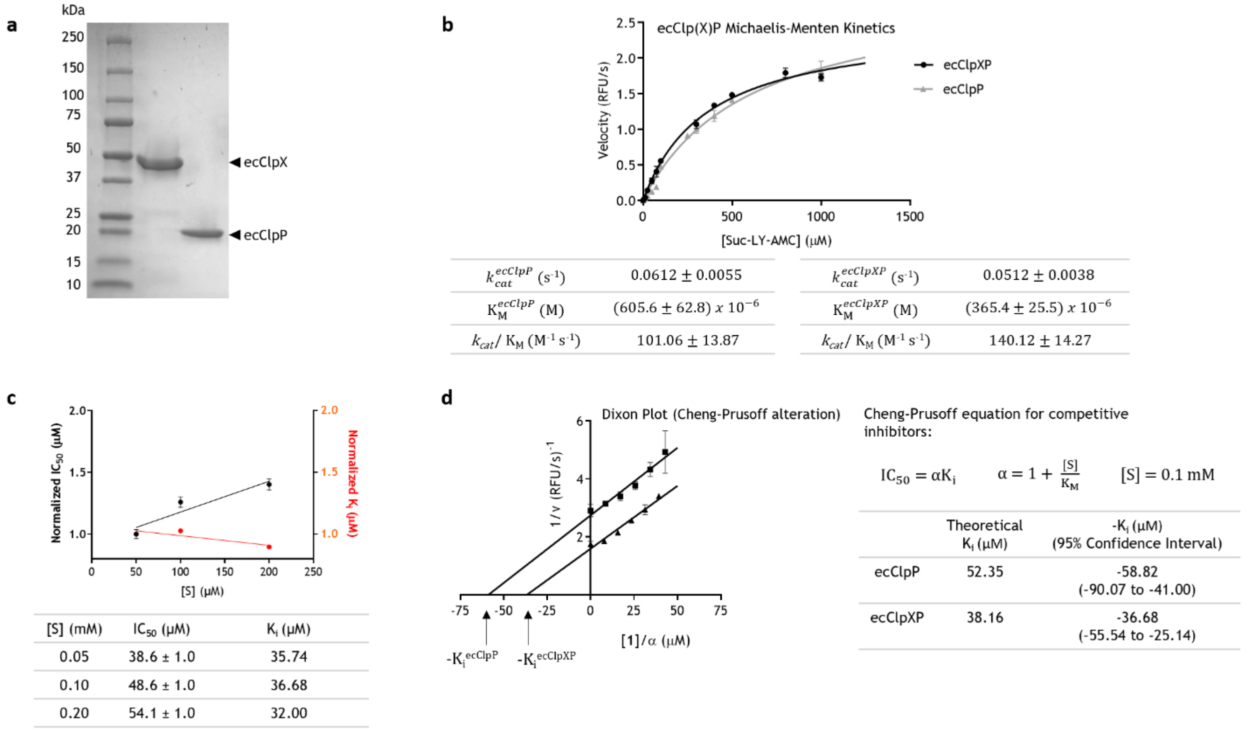


| | | | |
|---|---|---|---|
| $k_{cat}^{ecClpP}$ ($s^{-1}$) | $0.0612 \pm 0.0055$ | $k_{cat}^{ecClpXP}$ ($s^{-1}$) | $0.0512 \pm 0.0038$ |
| $K_M^{ecClpP}$ (M) | $(605.6 \pm 62.8)\ x\ 10^{-6}$ | $K_M^{ecClpXP}$ (M) | $(365.4 \pm 25.5)\ x\ 10^{-6}$ |
| $k_{cat}/K_M$ ($M^{-1} s^{-1}$) | $101.06 \pm 13.87$ | $k_{cat}/K_M$ ($M^{-1} s^{-1}$) | $140.12 \pm 14.27$ |

| [S] (mM) | $IC_{50}$ (μM) | $K_i$ (μM) |
|---|---|---|
| 0.05 | 38.6 ± 1.0 | 35.74 |
| 0.10 | 48.6 ± 1.0 | 36.68 |
| 0.20 | 54.1 ± 1.0 | 32.00 |

$IC_{50} = \alpha K_i \qquad \alpha = 1 + \frac{[S]}{K_M} \qquad [S] = 0.1\ mM$

| | Theoretical $K_i$ (μM) | $-K_i$ (μM) (95% Confidence Interval) |
|---|---|---|
| ecClpP | 52.35 | -58.82 (-90.07 to -41.00) |
| ecClpXP | 38.16 | -36.68 (-55.54 to -25.14) |

**Supplementary Figure 2.** Purification and kinetic characterization of ecClpXP inhibition. (a) SDS-PAGE of ecClpX and ecClpP, stained with Coomassie dye. (b) Michaelis-Menten curve of ecClpP and ecClpXP proteolysis. (c) Normalized Cheng-Prusoff variables showing an increase of $IC_{50}$ and stable $K_i$ with increasing substrate concentrations, supporting a competitive model of ecClpXP inhibition. (d) Cheng-Prusoff alteration of a Dixon plot to identify $K_i$ for ecClpP and ecClpXP. All kinetic curves were obtained in triplicates (n=3).

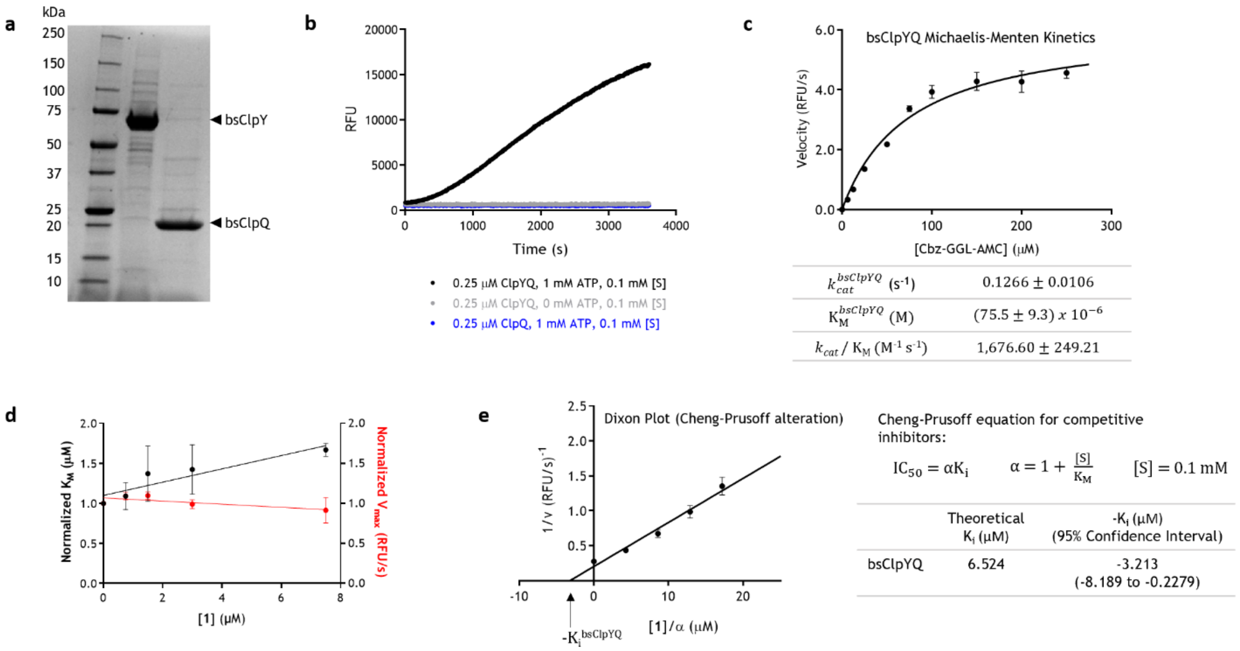


**Supplementary Figure 3.** Purification and kinetic characterization of bsClpYQ inhibition. (a) SDS-PAGE of bsClpY and bsClpQ, stained with Coomassie dye. (b) ATP- and ClpY- dependence of bsClpYQ proteolysis activity. (c) Michaelis-Menten curve of ClpYQ proteolysis. (d) Normalized competitive inhibition plot showing an increase of $K_M$ and stable $V_{max}$ with increasing inhibitor concentrations. (e) Cheng-Prusoff alteration of a Dixon plot to identify $K_i$. All kinetic curves were obtained in triplicates (n=3).

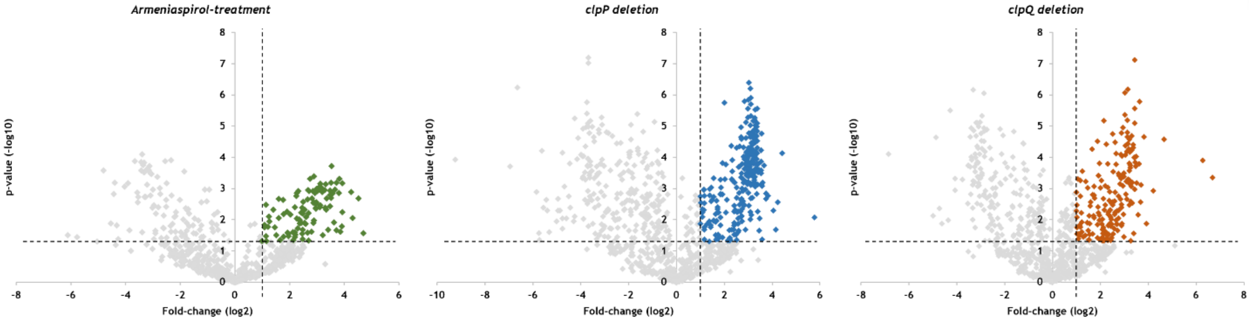


**Supplementary Figure 4.** Volcano plots of dimethyl labeling quantitative proteomics in *B. subtilis* of armeniaspirol, *ΔclpP* and *ΔclpQ* treatments with wild-type *B. subtilis* used as the control. Proteins of biological and statistical significance are highlighted in colour (log2 fold-change >2, p-value <0.05)

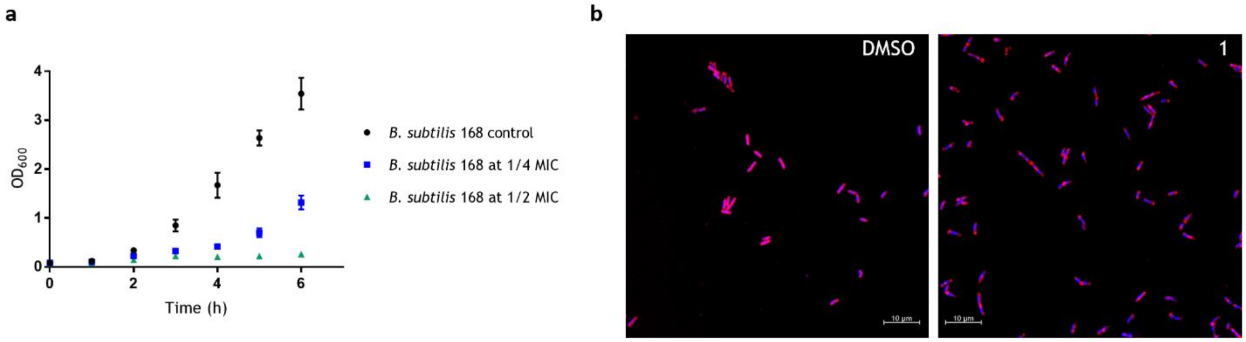


**Supplementary Figure 5.** Evaluation of phenotype in the absence and presence of armeniaspirol. (a) Growth curve of *B. subtilis* at sub-inhibitory concentrations of **1**. (b) Representative field-of-view of confocal fluorescent microscopy images of untreated (left) and treated (right) *B. subtilis* 168 at half the MIC (1 µg/ml). The nucleoid was stained with DAPI, while the membrane was stained with FM-64FX dye.

**Supplementary Table 1.** Comparison of armeniaspirol-treated proteome in relation to protein markers involved in various antibiotic treatments. Protein fold-change and p-values are identified for specific proteins associated with certain antibiotic treatments and are shown in tabular format. Each protein marker highlighted has been shown to have substantial enrichment with its corresponding antibiotic treatment[1-5]. Data analysis was performed as described in the Methods.

| Accession | Description | Fold-change | p-value |
|---|---|---|---|
| **Protein Biosynthesis Biomarkers** [ref: tetracycline, chelocardin] [1] | | | |
| P21464 | 30S ribosomal protein S2, RpsB | 4.101 | 0.007 |
| P21468 | 30S ribosomal protein S6, RpsF | 7.062 | 0.002 |
| P42923 | 50S ribosomal protein L10, RplJ | 0.646 | 0.555 |
| P80868 | Elongation factor G, FusA | 1.426 | 0.098 |
| P33166 | Elongation factor Tu, TufA | 0.906 | 0.767 |
| **Fatty Acid Biosynthesis Biomarkers** [ref: platencin, platensimycin, cerulenin, triclosan] [2] | | | |
| O34746 | 3-oxoacyl-[acyl-carrier-protein] synthase 3 protein 1, FabHA | 0.614 | 0.412 |
| O34340 | 3-oxoacyl-[acyl-carrier-protein] synthase 2, FabF | 0.742 | 0.587 |
| P54616 | Enoyl-[acyl-carrier-protein] reductase [NADH], FabI | 0.254 | 0.025 |
| **Cell Envelope Stress Biomarkers** [ref: merscadin, bacitracin, vancomycin] [3] | | | |
| P81100 | Stress response protein SCP2, YceC | 1.320 | 0.163 |
| O34833 | Uncharacterized protein YceH | 0.572 | 0.436 |
| **DNA Damage Response Biomarkers** [ref: mitomycin, daunomycin, Adriamycin] [4,5] | | | |
| P16971 | Protein RecA | 0.832 | 0.526 |

**Supplementary Table 2.** Checkerboard assays evaluating synergy. 5-chloro-armeniaspirol was paired with Gram-positive antibiotics of varying mechanisms in *B. subtilis*. The fractional inhibitory concentration (FIC) index was calculated according to the equation below, where $MIC_A$ represents the MIC of **1** alone, $MIC_{AC}$ represents the MIC of **1** in combination with antibiotic X, $MIC_X$ represents the MIC of antibiotic X alone, $MIC_{XC}$ represents the MIC of antibiotic X in combination with **1**. Synergy is defined as FIC index ≤ 0.5. No synergy is observed with **1**.

$$FIC = \frac{MIC_{AC}}{MIC_A} + \frac{MIC_{XC}}{MIC_X}$$

| Antibiotic X = | Tetracycline | Ciprofloxacin | Penicillin | Cerulenin |
|---|---|---|---|---|
| Mechanism of Action: | Protein synthesis | DNA replication | Cell wall biosynthesis | Fatty acid biosynthesis |
| | FIC index: | | | |
| 5-chloro-armeniaspirol, **1** | 0.625 | ≥ 1.000 | ≥ 1.000 | ≥ 1.000 |
| **Effect:** | **Indifferent** | **Indifferent** | **Indifferent** | **Indifferent** |

**Supplementary Table 3.** Enrichment of divisome- and elongasome-related proteins in **1**-treated, *ΔclpP*, and *ΔclpQ B. subtilis*, supporting a mechanism involving inhibition of ClpXP. Protein fold-change and p-values are provided (*n.d.*; not detected). Significance is defined as fold-change > 2 and p-value < 0.05 (*not significant, detected in 2 out of 3 replicates). Data analysis was performed as described in the Methods.

| **Divisome Proteins** | | **1-treatment** | | ***clpP* deletion** | | ***clpQ* deletion** | |
|---|---|---|---|---|---|---|---|
| *Accession* | *Description* | *fold-change* | *p-value* | *fold-change* | *p-value* | *fold-change* | *p-value* |
| P17865 | Cell division protein, FtsZ | 9.351 | 0.0312 | 8.324 | 0.0002 | 1.796 | 0.0150 |
| P28598 | 60 kDa chaperonin, GroEL | 2.860 | 0.0303 | 2.906 | 0.0005 | 1.085 | 0.5113* |
| P17820 | Chaperone protein, DnaK | 4.795 | 0.0014 | 2.744 | 0.0994 | 1.067 | 0.8579* |
| P80698 | Trigger factor, Tig | 11.539 | 0.0020 | 3.802 | 0.0002 | 0.797 | 0.3494* |
| O34894 | Septation ring formation regulator, EzrA | 0.043 | 0.0130 | 6.815 | 0.0031 | 0.497 | 0.1519* |
| P71021 | Septum site-determining protein, DivIVA | 7.360 | 0.0029 | 4.584 | 0.1126* | 4.792 | 0.0397 |
| P0CI74 | Cell cycle protein, GpsB | 3.903 | 0.1247* | *n.d.* | | *n.d.* | |
| Q01464 | Septum site-determining protein, MinD | 3.686 | 0.1711* | 8.070 | 4.9180E-05 | *n.d.* | |
| **Elongasome Proteins** | | **1-treatment** | | ***clpP* deletion** | | ***clpQ* deletion** | |
| *Accession* | *Description* | *fold-change* | *p-value* | *fold-change* | *p-value* | *fold-change* | *p-value* |
| Q01465 | Rod shape-determining protein, MreB | 5.123 | 0.0098 | 6.052 | 0.0002 | 1.282 | 0.1680* |
| P39751 | MreB-like protein, Mbl | *n.d.* | | 16.574 | 0.0052 | 14.242 | 2.1970E-05 |
| O32023 | Uncharacterized protein, YqzC | 11.245 | 0.0016 | 8.023 | 0.0014 | *n.d.* | |

**Supplementary Table 4.** List of *Bacillus subtilis* 168 gene deletion strains used in study.

| [a]**BGSC №** | **Original Strain** | **Genotype** | **PMID Ref.** | **MIC (µg/ml)** |
|---|---|---|---|---|
| BKE16150 | Bacillus subtilis 168 | ΔclpQ::erm trpC2 | 28189581 | 2 |
| BKK34540 | Bacillus subtilis 168 | ΔclpP::kan trpC2 | 28189581 | 2 |
| [b]**NBRP** | **Original Strain** | **Genotype** | **PMID Ref.** | **MIC (µg/ml)** |
| MGNA-A086 | Bacillus subtilis 168 | ΔclpY::pMUTIN (erm) | 12682299 | 2 |

[a]BGSC, Bacillus Genetic Stock Center, Columbus, OH
[b]NBRP, National BioResource Project: B. subtilis, NIG, Japan

**Supplementary Table 5**. List of plasmids used in study. All plasmids were heterologously expressed in *Escherichia coli* BL21 (DE3).

| **Plasmid** | **Inserted Gene** | **Antibiotic Marker** | **Parental Vector** |
|---|---|---|---|
| pPL29 | *clpQ* (*B. subtilis* 168) | $Amp^R$ | pET21c |
| pPL31 | *clpY* (*B. subtilis* 168) | $Kan^R$ | pET28b |
| pProEx-Htb-ClpX | *clpX* (*E. coli*) | $Amp^R$ | pProEx Htb |
| pET9a-ClpP | *clpP* (*E. coli*) | $Kan^R$ | pET9a |

### Synthetic methods and characterization

#### General information

All reactions were performed in oven-dried or flame-dried glass round-bottom flasks equipped with magnetic stir bars. Purification of reaction products was carried out by flash column chromatography using silica gel. $^{1}H$ NMR spectra were recorded on either a Bruker 400 MHz or Bruker 300 MHz at ambient temperature. $^{13}C$ NMR spectra were recorded on either a Bruker 100 MHz or Bruker 75 MHz at ambient temperature. Spectra are recorded in parts per million using residual solvent as the internal standard ($(CD_3)_2SO$ at 2.50 ppm, $CDCl_3$ at 7.26 ppm and $CD_3OD$ at 3.31 ppm for 1H NMR and $(CD_3)_2SO$ at 39.52 ppm, $CDCl_3$ at 77.16 ppm and $CD_3OD$ at 49.00 ppm for $^{13}C$ NMR.) $^{1}H$ NMR data was reported as: multiplicity (br = broad, s = singlet, d = doublet, t = triplet, q = quartet, quin. = quintet, sext. = sextuplet, sept = septuplet, m = multiplet), integration and coupling constant(s) in Hz. High-resolution mass spectrometry was performed by electrospray ionization (ESI) in positive ion or negative ion mode using a Micromass Q-TOF 1 mass spectrometer.

#### Materials

Unless otherwise noted, all commercially available materials were purchased from commercial sources and used without further purification.

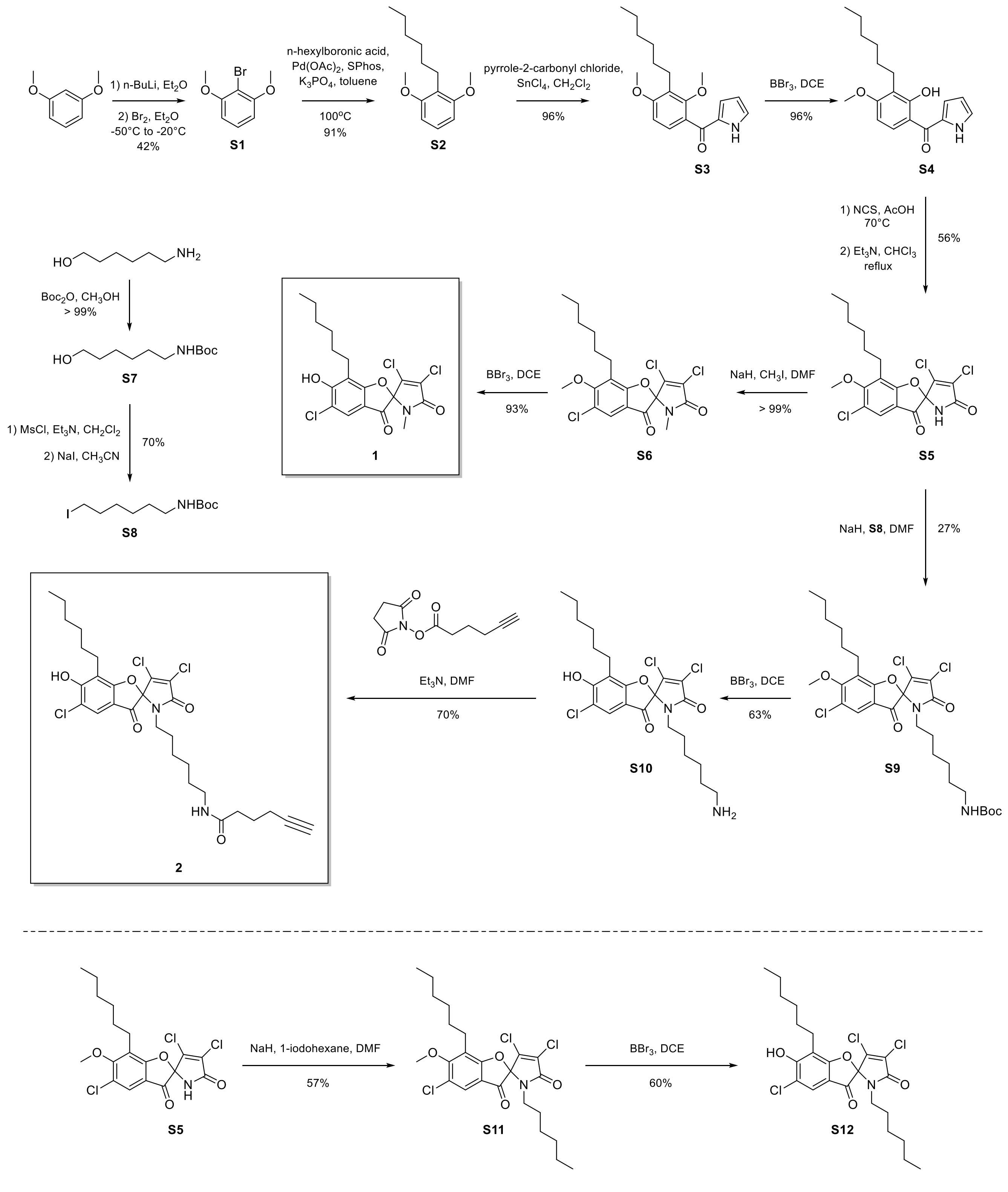


**Supplementary Scheme 1.** Synthesis of 5-chloro-armeniaspirol A and analogs

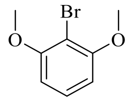

**1-Bromo-2,6-dimethoxybenzene**, (**S1**)
1,3-dimethoxybenzene (14.40 g, 0.104 mol, 1.0 equiv.) was dissolved in $Et_2O$ (375 mL) in a round-bottom flask. n-BuLi (50 mL of a 2.5 M solution, 1.2 equiv.) was added and the solution was allowed to stir at ambient temperature for 4 hours. The reaction mixture was then cooled to – 50 °C using a dry ice/acetone bath. Bromine (18.65 g, 0.117 mol, 1.1 equiv.) was added dropwise. The solution was then heated to room temperature and allowed to react for another two hours. To the resulting mixture, 250 mL of a 10 % sodium thiosulfate solution was added and the resulting mixture was allowed to stir for 1 hour. The solution was extracted 2 × with $Et_2O$. The resulting organic fractions were combined and washed with brine. The organic phase was then dried with $Na_2SO_4$ and concentrated to yield the desired compound (9.45 g, 43.5 mmol, 42 % yield) which was used without further purification. The NMR data were consistent with literature values[6]. $^1$H NMR (400 MHz, $CDCl_3$) δ 7.23 (t, *J* = 8.3 Hz, 1H), 6.58 (d, *J* = 8.4 Hz, 1H), 3.90 (s, 3H). $^{13}$C NMR (100 MHz, $CDCl_3$) δ 157.34, 128.38, 104.82, 101.08, 56.58.

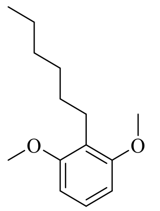

**2-Hexyl-1,3-dimethoxy-benzene**, (**S2**)
In a round-bottom flask, **S1** (5.0 g, 23.0 mmol, 1.0 equiv.) was dissolved in toluene (230 mL). To the resulting solution, *n*-hexylboronic acid (5.98 g, 46 mmol, 2.0 equiv.), $K_3PO_4{\cdot}H_2O$ (10.6 g, 46 mmol, 2.0 equiv.), $Pd(OAc)_2$ (516 mg, 2.3 mmol, 0.1 equiv.), and SPhos (1.89 g, 4.6 mmol, 0.2 equiv.) were added at room temperature. The mixture was stirred at 100 °C for 15 hours. After cooling, the reaction mixture was quenched with $NH_4Cl_{(aq)}$ and extracted 3 × with EtOAc. The organic fractions were combined, washed with brine, dried over $Na_2SO_4$ and concentrated. The desired compound (4.65 g, 20.9 mmol, 91 % yield) was purified from the crude mixture by silica column chromatography (100 % hexanes). The NMR data were consistent with literature values[7]. $^1$H NMR (300 MHz, DMSO) δ 7.10 (t, *J* = 8.3 Hz, 1H), 6.59 (d, *J* = 8.3 Hz, 2H), 3.74 (s, 6H), 2.56 – 2.51 (m, 2H), 1.43 – 1.31 (m, 2H), 1.29 – 1.19 (m, 6H), 0.85 (d, *J* = 6.9 Hz, 3H). $^{13}$C NMR (75 MHz, DMSO) δ 157.75, 126.78, 117.98, 103.85, 55.59, 31.17, 28.82, 28.81, 22.38, 22.11, 13.98.

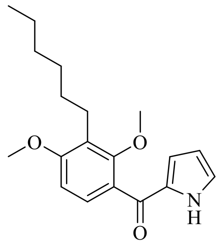

**(3-Hexyl-2,4-dimethoxy-phenyl)-(1*H*-pyrrol-2-yl)-methanone**, (**S3**)
In a round-bottom flask, **S2** (4.09 g, 18.4 mmol, 1.0 equiv.) was dissolved in $CH_2Cl_2$ (92 mL) and cooled to 0 °C with an ice bath. 1H-Pyrrole-2-carbonyl chloride (46 mmol, 2.5 equiv.) was added, followed by $SnCl_4$ (92 mL of a 1.0 M solution, 5 equiv.). The mixture was stirred for 1 hour at 0 °C, then warmed to ambient temperature. The reaction mixture was quenched with a saturated $NaHCO_{3(aq)}$ and extracted 3 × with EtOAc. The organic fractions were combined, washed with brine, dried over $Na_2SO_4$ and concentrated. The desired compound (5.59 g, 17.7 mmol, 96 % yield) was purified from the crude mixture by silica column chromatography (10 to 25 % EtOAc in hexanes). The NMR data were consistent with literature values[7]. $^1$H NMR (400 MHz, DMSO) δ 11.98 (s, 1H), 7.25 (d, *J* = 8.5 Hz, 1H), 7.18 – 7.13 (m, 1H), 6.80 (d, *J* = 8.6 Hz, 1H), 6.51 – 6.48 (m, 1H), 6.18 (dt, *J* = 4.2, 2.3 Hz, 1H), 3.83 (s, 3H), 3.59 (s, 3H), 2.61 – 2.54 (m, 2H), 1.51 – 1.40 (m, 2H), 1.35 – 1.22 (m, 6H), 0.86 (t, *J* = 6.8 Hz, 3H). $^{13}$C NMR (100 MHz, DMSO) δ 183.86, 159.98, 157.39, 132.47, 128.49, 126.58, 125.90, 124.13, 119.54, 110.44, 105.88, 62.55, 56.26, 31.54, 29.64, 29.43, 23.61, 22.52, 14.41.

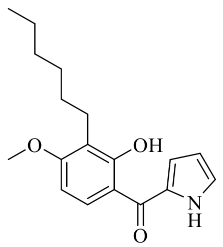

**(3-Hexyl-2-hydroxy-4-methoxy-phenyl)-(1*H*-pyrrol-2-yl)-methanone**, (**S4**)
In a round-bottom flask, **S3** (5.585 g, 17.7 mmol, 1.0 equiv.) was dissolved in 1,2-dichloroethane (21 mL). The solution was cooled to -20 °C using a dry ice/acetone bath. $BBr_3$ (21.2 mL of a 1 M solution, 1.2 equiv.) was added dropwise and the reaction mixture was stirred at -20 to – 10 °C for 2 hours. Subsequently, $Et_3N$ / water was added and the solution was extracted 3 × with EtOAc. The organic fractions were combined, washed with brine, dried over $Na_2SO_4$, and concentrated. The desired compound (5.15 g, 17.1 mmol, 97 % yield) was purified from the crude mixture by silica column chromatography (15 % EtOAc in hexanes). The NMR data were consistent with literature values[7]. $^1H$ NMR (400 MHz, DMSO) δ 12.08 (s, 1H), 7.95 (d, *J* = 9.0 Hz, 1H), 7.25 – 7.22 (m, 1H), 6.99 (ddd, *J* = 3.8, 2.4, 1.3 Hz, 1H), 6.66 (d, *J* = 9.1 Hz, 1H), 6.31 (dt, *J* = 4.0, 2.3 Hz, 1H), 3.86 (s, 3H), 3.35 (s, 3H), 2.62 – 2.54 (m, 2H), 1.49 – 1.38 (m, 2H), 1.33 – 1.21 (m, 6H), 0.84 (t, *J* = 6.9 Hz, 3H). $^{13}C$ NMR (100 MHz, DMSO) δ 185.75, 162.24, 160.99, 130.71, 129.36, 126.34, 119.05, 117.12, 113.09, 110.73, 102.60, 55.85, 31.17, 28.86, 28.30, 22.10, 22.08, 13.96.

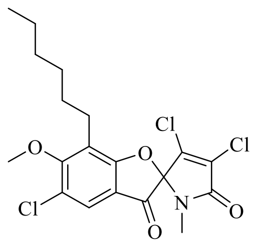

**3',4',5-Trichloro-7-hexyl-6-methoxy-1'-methyl-1'*H*-spiro[1-benzofuran-2,2'-pyrrole]-3,5'-dione**, (**S6**)
In a round-bottom flask, **S4** (1.50 g, 5.0 mmol, 1.0 equiv.) was dissolved in acetic acid (50 mL). *N*-chlorosuccinimide (1.33 g, 10.0 mmol, 2.0 equiv.) was added and the mixture was stirred at ambient temperature for 2 hours. Following this, *N*-chlorosuccinimide (2.67 g, 20 mmol, 4.0 equiv.) was added and the resulting mixture was heated to 70 °C for 16 hours. The reaction mixture was then quenched with a 10 % $K_2CO_{3(aq)}$ and extracted 3 × with EtOAc. The organic fractions were combined, washed with brine, dried over $Na_2SO_4$ and concentrated. The resulting oil was dissolved in $CHCl_3$ (33 mL) and $Et_3N$ (1.4 mL) was added. The mixture was heated at 60 °C for 5 hours. The solution was cooled to ambient temperature, concentrated, and the spiro- intermediate 5,3',4'-trichloro-6-methoxy-7-hexyl-spiro[benzofuran-2(3H),2'-[2H]pyrrole]-3,5'(1'H)-dione (S5) (1.133 g, 2.8 mmol, 56 % yield over two steps) was isolated by from the crude mixture by silica column chromatography (2 % to 20 % EtOAc in hexanes). The *spiro*-intermediate (40 mg, 0.096 mmol, 1.0 equiv.) was dissolved in DMF (2 mL). The solution was cooled to 0 °C using an ice bath and NaH (3.4 mg, 0.143 mmol, 1.5 equiv.) was added and allowed to stir for 15 min. Subsequently, MeI (17.6 mg, 0.124 mmol, 1.3 equiv.) was added dropwise and stirring was continued at 0 °C to ambient temperature for 5 hours. The mixture was quenched with $NH_4Cl_{(aq)}$, extracted 3 × with EtOAc, washed with brine, dried over $Na_2SO_4$, and concentrated. The desired compound (40.0 mg, 0.095 mmol, >99% yield) was purified from the crude mixture by silica column chromatography (15 % EtOAc in hexanes). The NMR data were consistent with literature values[7]. $^1H$ NMR (400 MHz, $CDCl_3$) δ 7.63 (s, 1H), 4.00 (s, 3H), 2.79 (s, 3H), 2.76 – 2.70 (m, 2H), 1.64 – 1.55 (m, 3H), 1.41 – 1.24 (m, 6H), 0.88 (t, *J* = 7.0 Hz, 3H). $^{13}C$ NMR (100 MHz, $CDCl_3$) δ 189.88, 170.33, 163.79, 163.16, 138.24, 129.38, 124.84, 124.33, 123.73, 115.65, 97.05, 61.92, 31.73, 29.46, 29.38, 25.96, 24.00, 22.71, 14.19. HRMS (ESI): Exact mass calculated for $C_{19}H_{20}Cl_3NNaO_4$ $[M + Na]^+$: 454.0350. Found: 454.0356

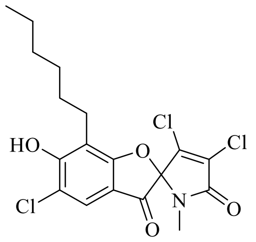

**3',4',5-Trichloro-7-hexyl-6-hydroxy-1'-methyl-1'*H*-spiro[1-benzofuran-2,2'-pyrrole]-3,5'-dione**, (**1**)

In a round-bottom flask, S6 (54.2 mg, 0.125 mmol, 1.0 equiv.) was dissolved in 1,2-dichloroethane (1 mL). The mixture was cooled to 0 °C using an ice bath and $BBr_3$ (380 µL of a 1 M solution, 3.0 equiv.) was added dropwise. The reaction was allowed to proceed for 4 hours from 0 °C to ambient temperature. $Et_3N$ / water was added and the solution was extracted 3 × with EtOAc. The organic fractions were combined, washed with brine, dried over $Na_2SO_4$, and concentrated. The desired compound (48.7 mg, 0.116 mmol, 93 %) was purified from the crude mixture by silica column chromatography (5 to 20 % EtOAc in hexanes). The NMR data were consistent with literature values[7]. $^1H$ NMR (400 MHz, $CDCl_3$) δ 7.63 (s, 1H), 6.62 (s, 1H), 2.79 (s, 3H), 2.76 (t, *J* = 7.6 Hz, 2H), 1.66 – 1.56 (m, 2H), 1.39 – 1.24 (m, 6H), 0.87 (t, *J* = 7.0 Hz, 3H). $^{13}C$ NMR (100 MHz, $CDCl_3$) δ 188.93, 170.34, 163.21, 158.77, 138.43, 129.27, 122.56, 117.67, 115.79, 112.66, 97.16, 31.77, 29.22, 28.47, 25.91, 23.37, 22.74, 14.21. HRMS (ESI): Exact mass calculated for $C_{18}H_{17}Cl_3NO_4$ $[M - H]^-$: 416.0229. Found: 416.0223

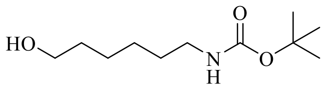

**6-(*tert*-Butoxycarbonylamino)-1-hexanol**, (**S7**)

In a round-bottom flask, 6-amino-1-hexanol (2.0 g, 17.1 mmol, 1.0 equiv.) was dissolved in methanol (40 mL). Di-tert-butyl dicarbonate (4.1 g, 18.8 mmol, 1.1 equiv.) was added and the reaction was stirred for 1.5 hours at ambient temperature. The reaction mixture was concentrated and the desired product (3.72 g, 17.1 mmol, >99 % yield) was purified from the crude residue by silica column chromatography (2 % methanol in $CH_2Cl_2$). The NMR data were consistent with literature values[8]. $^1H$ NMR (300 MHz, $CDCl_3$) δ 3.63 (t, *J* = 6.5 Hz, 2H), 3.11 (t, *J* = 6.9 Hz, 2H), 1.62 – 1.29 (m, 17H). $^{13}C$ NMR (75 MHz, $CDCl_3$) δ 156.21, 79.21, 62.71, 40.50, 32.68, 30.16, 28.52, 26.50, 25.39.

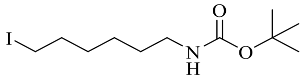

**(6-Iodo-hexyl)-carbamic acid tert-butyl ester**, (**S8**)

S7 (2.25 g, 10.35 mmol, 1.0 equiv.) was dissolved in $CH_2Cl_2$ (30 mL) in a round-bottom flask. $Et_3N$ (2.2 mL, 15.53 mmol, 1.5 equiv.) was added and the solution was cooled to 0 °C using an ice bath. Methanesulfonyl chloride (881 µL, 11.39 mmol, 1.1 equiv.) was added dropwise and the mixture was stirred for 2 hours, while warming to ambient temperature. The resulting solution was washed with $H_2O$, washed with brine, dried over $Na_2SO_4$ and concentrated. The crude oil was dissolved in acetonitrile (45 mL) in a round-bottom flask and NaI (7.73 g, 51.6 mmol, 5 equiv.) was added. The mixture was stirred at room temperature for 16 hours. The reaction mixture was concentrated, and the residue was partitioned between EtOAc and $H_2O$. The organic phase was separated, washed with brine, dried over $Na_2SO_4$ and concentrated. The desired compound (2.37 g, 7.24 mmol, 70 % yield) was purified from the crude mixture by silica column chromatography (10 % EtOAc in hexanes). The NMR data were consistent with literature values[8]. $^1H$ NMR (400 MHz, $CDCl_3$) δ 4.52 (br, 1H), 3.17 (t, *J* = 7.0 Hz, 2H), 3.13 – 3.05 (m, 2H), 1.86 – 1.75 (m, 2H), 1.53 – 1.27 (m, 14H). $^{13}C$ NMR (100 MHz, $CDCl_3$) δ 156.10, 79.20, 40.60, 33.49, 30.26, 30.04, 28.55, 25.84, 7.05.

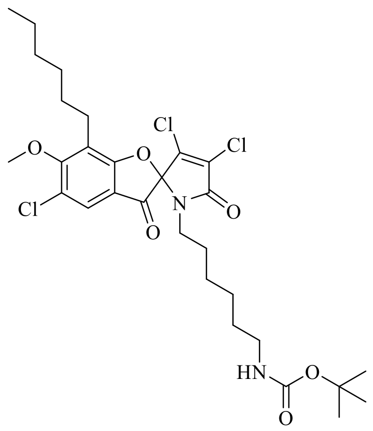

**3',4',5-Trichloro-7-hexyl-6-methoxy-1'-[6-(*tert*-butoxycarbonylamino)hexyl]-1'*H*-spiro[1-benzofuran-2,2'-pyrrole]-3,5'-dione**, (**S9**)

The *spiro*- intermediate **S5** (80 mg, 0.191 mmol, 1.0 equiv.) was dissolved in DMF (2 mL). The solution was cooled to 0 °C using an ice bath and NaH (5.95 mg, 0.248 mmol, 1.3 equiv.) was added and allowed to stir for 15 min. Subsequently, (6-iodo-hexyl)-carbamic acid tert-butyl ester (75 mg, 0.229 mmol, 1.2 equiv.) was added and stirring was continued at 0 °C to ambient temperature for 5 hours. The mixture was quenched with $NH_4Cl_{(aq)}$, extracted 3 × with EtOAc, washed with brine, dried over $Na_2SO_4$, and concentrated. The desired compound (32.4 mg, 0.053 mmol, 27 % yield) was purified from the crude mixture by silica column chromatography (0 to 15 % EtOAc in hexanes). $^1$H NMR (400 MHz, $CDCl_3$) δ 7.63 (s, 1H), 4.50 (br, 1H), 4.00 (s, *J* = 9.0 Hz, 3H), 3.37 (dt, *J* = 14.6, 7.3 Hz, 1H), 3.13 – 2.97 (m, 3H), 2.79 – 2.63 (m, 2H), 1.65 – 1.17 (m, 28H), 0.88 (s, 3H). $^{13}$C NMR (100 MHz, $CDCl_3$) δ 190.21, 169.97, 163.75, 163.50, 156.09, 138.31, 129.24, 124.80, 124.30, 123.75, 115.77, 97.28, 61.94, 41.40, 40.53, 31.73, 29.98, 29.83, 29.51, 29.42, 28.71, 28.55, 26.41, 26.31, 23.98, 22.69, 14.20. HRMS (ESI): Exact mass calculated for $C_{29}H_{39}Cl_3N_2NaO_6$ $[M + H]^+$: 639.1766. Found: 639.1771

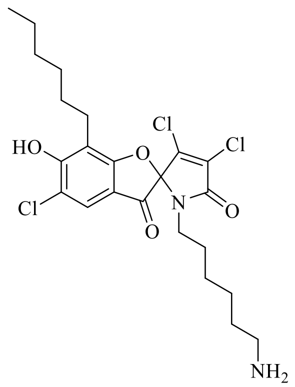

**1'-(6-Aminohexyl)-3',4',5-trichloro-7-hexyl-6-hydroxy-1'*H*-spiro[1-benzofuran-2,2'-pyrrole]-3,5'-dione**, (**S10**)

In a round-bottom flask, **S9** (30 mg, 0.0485 mmol, 1.0 equiv.) was dissolved in 1,2-dichloroethane (1 mL). The mixture was cooled to 0 °C using an ice bath and $BBr_3$ (243 µL of a 1 M solution, 5.0 equiv.) was added dropwise. The reaction was allowed to proceed for 4 hours from 0 °C to ambient temperature. $Et_3N$ / water was added and the solution was extracted 3 × with EtOAc. The organic fractions were combined, washed with brine, dried over $Na_2SO_4$, and concentrated. The desired compound (15.4 mg, 0.0305 mmol, 63 %) was purified from the crude mixture by silica column chromatography (5 to 20 % methanol in $CH_2Cl_2$). $^1$H NMR (400 MHz, MeOD) δ 7.44 (s, 1H), 3.41 (dt, *J* = 14.3, 7.1 Hz, 1H), 3.35 (s, 1H), 3.09 (dt, *J* = 14.4, 7.1 Hz, 1H), 2.85 (t, *J* = 7.5 Hz, 2H), 2.65 – 2.49 (m, 2H), 1.62 – 1.46 (m, 6H), 1.40 – 1.23 (m, 10H), 0.88 (t, *J* = 6.6 Hz, 3H). $^{13}$C NMR (100 MHz, MeOD) δ 185.04, 176.36, 171.68, 165.19, 142.00, 128.54, 126.71, 122.87, 115.03, 104.80, 99.00, 41.68, 40.60, 33.07, 30.43, 29.85, 29.25, 28.33, 27.09, 26.71, 24.16, 23.75, 14.50.

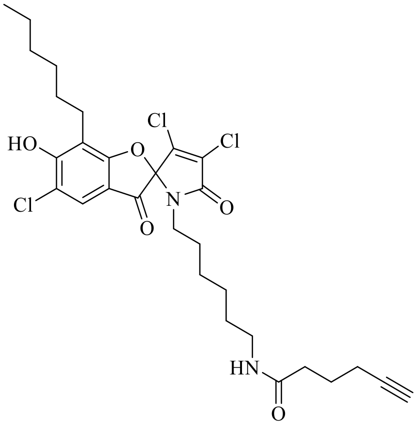


**3',4',5-Trichloro-7-hexyl-1'-[6-(5-hexynoylamino)hexyl]-6-hydroxy-1'*H*-spiro[1-benzofuran-2,2'-pyrrole]-3,5'-dione**, (**2**)

In a round-bottom flask, **S10** (87.8 mg, 0.175 mmol, 1.0 equiv.) was dissolved in DMF (1.75 mL). 2,5-Dioxo-1-pyrrolidinyl 5-hexynoate (37 mg, 0.175, 1.0 equiv.) and $Et_3N$ (12 µl, 0.0875 mmol, 0.5 equiv.) were added to the mixture and the solution was stirred at room temperature for 6 hours. The reaction mixture was quenched with $NH_4Cl_{(aq)}$, extracted with EtOAc, washed with brine, dried over $Na_2SO_4$ and concentrated. The desired product (72.8 mg, 0.122 mmol, 70 % yield) was purified by silica column chromatography (5 to 25 % EtOAc in hexanes). $^1$H NMR (400 MHz, MeOD) δ 7.67 (s, 1H), 3.44 (dt, *J* = 14.5, 7.2 Hz, 1H), 3.11 (t, *J* = 7.0 Hz, 2H), 3.08 – 3.01 (m, 1H), 2.81 – 2.68 (m, 2H), 2.28 (d, *J* = 7.3 Hz, 2H), 2.25 – 2.17 (m, 3H), 1.78 (p, *J* = 7.1 Hz, 2H), 1.67 – 1.56 (m, 2H), 1.51 – 1.22 (m, 14H), 0.89 (t, *J* = 6.7 Hz, 3H). $^{13}$C NMR (101 MHz, MeOD) δ 189.88, 175.24, 171.17, 164.73, 163.07, 140.26, 129.59, 124.10, 120.14, 117.01, 112.69, 98.52, 84.12, 70.20, 49.85, 42.19, 40.19, 35.81, 32.81, 30.14, 29.59, 29.52, 27.36, 27.33, 25.97, 23.98, 23.64, 18.64, 14.44. HRMS (ESI): Exact mass calculated for $C_{29}H_{35}Cl_3N_2NaO_5$ $[M + Na]^+$: 619.1504. Found: 619.1509.

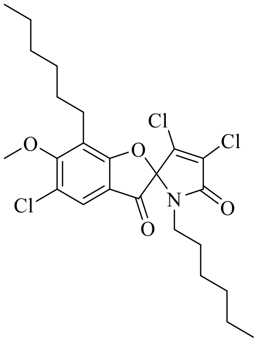


**3',4',5-Trichloro-7-hexyl-1'-hexyl-6-methoxy-1'*H*-spiro[1-benzofuran-2,2'-pyrrole]-3,5'-dione**, (**S11**)

The *spiro*- intermediate **S5** (360 mg, 0.936 mmol, 1.0 equiv.) was dissolved in DMF (5 mL). The solution was cooled to 0 °C using an ice bath and NaH (48.8 mg, 1.22 mmol, 1.3 equiv.) was added and allowed to stir for 15 min. Subsequently, 1-iodohexane (238 mg, 1.123 mmol, 1.2 equiv.) was added dropwise and stirring was continued at 0 °C to ambient temperature for 5 hours. The mixture was quenched with $NH_4Cl_{(aq)}$, extracted 3 × with EtOAc, washed with brine, dried over $Na_2SO_4$, and concentrated. The desired compound (269 mg, 0.535 mmol, 57 % yield) was purified from the crude mixture by silica column chromatography (15 % EtOAc in hexanes). $^1$H NMR (400 MHz, $CDCl_3$) δ 7.63 (s, 1H), 4.00 (s, 3H), 3.42 (dt, *J* = 14.5, 7.3 Hz, 1H), 3.00 (dt, *J* = 14.5, 7.3 Hz, 1H), 2.78 – 2.64 (m, 2H), 1.65 – 1.53 (m, 2H), 1.48 – 1.12 (m, 14H), 0.88 (t, *J* = 6.9 Hz, 3H), 0.83 (t, *J* = 6.8 Hz, 3H). $^{13}$C NMR (100 MHz, $CDCl_3$) δ 190.25, 170.00, 163.71, 163.46, 138.25, 129.29, 124.75, 124.33, 123.72, 115.84, 97.38, 61.93, 41.67, 31.74, 31.36, 29.54, 29.47, 28.76, 26.47, 24.00, 22.71, 22.56, 14.20, 14.10. HRMS (ESI): Exact mass calculated for $C_{24}H_{30}Cl_3NNaO_4$ $[M + Na]^+$: 524.1133. Found: 524.1133

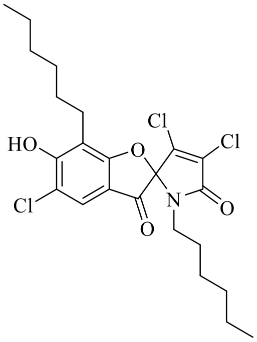


**3',4',5-Trichloro-7-hexyl-1'-hexyl-6-hydroxy-1'*H*-spiro[1-benzofuran-2,2'-pyrrole]-3,5'-dione**, (**S12**)
In a round-bottom flask, S11 (100 mg, 0.198 mmol, 1.0 equiv.) was dissolved in 1,2-dichloroethane (1.4 mL). The mixture was cooled to 0 °C using an ice bath and $BBr_3$ (596 µL of a 1 M solution, 3.0 equiv.) was added dropwise. The reaction was allowed to proceed for 4 hours from 0 °C to ambient temperature. $Et_3N$ / water was added and the solution was extracted 3 × with EtOAc. The organic fractions were combined, washed with brine, dried over $Na_2SO_4$, and concentrated. The desired compound (58.3 mg, 0.119 mmol, 60 %) was purified from the crude mixture by silica column chromatography (5 to 20 % EtOAc in hexanes). $^1H$ NMR (400 MHz, $CDCl_3$) δ 7.62 (s, 1H), 6.51 (s, 1H), 3.46 – 3.36 (m, 1H), 3.04 – 2.91 (m, 1H), 2.80 – 2.63 (m, 2H), 1.65 – 1.11 (m, 16H), 0.86 (t, *J* = 7.1 Hz, 3H), 0.81 (t, *J* = 6.9 Hz, 3H). $^{13}C$ NMR (100 MHz, $CDCl_3$) δ 189.33, 170.01, 163.50, 158.65, 138.42, 129.20, 122.54, 117.57, 115.82, 112.88, 97.50, 41.65, 31.79, 31.39, 29.31, 28.76, 28.56, 26.50, 23.38, 22.74, 22.58, 14.22, 14.10. HRMS (ESI): Exact mass calculated for $C_{23}H_{27}Cl_3NO_4$ $[M - H]^-$: 486.1011. Found: 486.1006

### References


(1) Stepanek, J. J.; Lukežič, T.; Teichert, I.; Petković, H.; Bandow, J. E. Dual mechanism of action of the atypical tetracycline chelocardin. *Biochim. Biophys. Acta - Proteins Proteomics* **2016**, *1864* (6), 645–654.

(2) Wenzel, M.; Patra, M.; Albrecht, D.; Chen, D. Y. K.; Nicolaou, K. C.; Metzler-Nolte, N.; Bandow, J. E. Proteomic signature of fatty acid biosynthesis inhibition available for in vivo mechanism-of-action studies. *Antimicrob. Agents Chemother.* **2011**, *55* (6), 2590–2596.

(3) Wenzel, M.; Kohl, B.; Münch, D.; Raatschen, N.; Albada, H. B.; Hamoen, L.; Metzler-Nolte, N.; Sahl, H.-G.; Bandow, J. E. Proteomic Response of Bacillus subtilis to Lantibiotics Reflects Differences in Interaction with the Cytoplasmic Membrane. *Antimicrob. Agents Chemother.* **2012**, *56* (11), 5749–5757.

(4) Bandow, J. E.; Brötz, H.; Leichert, L. I. O.; Labischinski, H.; Hecker, M. Proteomic approach to understanding antibiotic action. *Antimicrob. Agents Chemother.* **2003**, *47* (3), 948–955.

(5) Sender, U.; Bandow, J.; Engelmann, S.; Lindequist, U.; Hecker, M. Proteomic signatures for daunomycin and adriamycin in Bacillus subtilis. *Pharmazie* **2004**, *59* (1), 65–70.

(6) Kamikawa, T.; Hayashi, T. Enantioposition-selective arylation of biaryl ditriflates by palladium-catalyzed asymmetric Grignard cross-coupling. *Tetrahedron* **1999**, *55*, 3455–3466.

(7) Couturier, C.; Bauer, A.; Rey, A.; Schroif-Dufour, C.; Broenstrup, M. Armeniaspiroles, a new class of antibacterials: antibacterial activities and total synthesis of 5-chloro-Armeniaspirole A. *Bioorg. Med. Chem. Lett.* **2012**, *22* (19), 6292–6296.

(8) Isomura, S.; Wirsching, P.; Janda, K. D. An immunotherapeutic program for the treatment of nicotine addiction: hapten design and synthesis. *J. Org. Chem.* **2001**, *66* (12), 4115–4121.
